# Histone demethylase KDM2A is a selective vulnerability of cancers relying on alternative telomere maintenance

**DOI:** 10.1101/2023.02.10.528023

**Authors:** Fei Li, Yizhe Wang, Inah Hwang, Ja-Young Jang, Libo Xu, Zhong Deng, Eun Young Yu, Yiming Cai, Caizhi Wu, Zhenbo Han, Yu-Han Huang, Xiangao Huang, Ling Zhang, Jun Yao, Neal F. Lue, Paul M. Lieberman, Haoqiang Ying, Jihye Paik, Hongwu Zheng

## Abstract

Telomere length maintenance is essential for cellular immortalization and tumorigenesis. 5% - 10% of human cancers rely on a recombination-based mechanism termed alternative lengthening of telomeres (ALT) to sustain their replicative immortality, yet there are currently no targeted therapies. Through CRISPR/Cas9-based genetic screens in an ALT-immortalized isogenic cellular model, here we identify histone lysine demethylase KDM2A as a molecular vulnerability selectively for cells contingent on ALT-dependent telomere maintenance. Mechanistically, we demonstrate that KDM2A is required for dissolution of the ALT-specific telomere clusters following homology-directed telomere DNA synthesis. We show that KDM2A promotes de-clustering of ALT multitelomeres through facilitating isopeptidase SENP6-mediated SUMO deconjugation at telomeres. Inactivation of KDM2A or SENP6 impairs post-recombination telomere de-SUMOylation and thus dissolution of ALT telomere clusters, leading to gross chromosome missegregation and mitotic cell death. These findings together establish KDM2A as a selective molecular vulnerability and a promising drug target for ALT-dependent cancers.

## Introduction

Telomeres are specialized nucleoprotein structures that shield the linear chromosome ends of eukaryotes from promiscuous DNA repair and nucleolytic degradation activities ^1^. Due to the chromosome “end-replication” problem, telomeric DNA undergoes progressive attrition with each cell division ^2^. Consequentially, proliferative tumor cells necessitate counteracting activity to maintain adequate telomere length and sustain their replicative immortality. While a majority of human cancers achieve this through telomerase activation that adds *de novo* telomere repeats to chromosome ends, the remaining 5% - 10% of them rely on a homologous recombination-based mechanism termed alternative lengthening of telomeres (ALT) ^3, 4^. Recent studies further reveal ALT as a conservative DNA damage repair pathway analogous to break-induced replication (BIR) in budding yeast ^5-8^. Despite the advancement, the molecular pathway(s) that control ALT activation and termination remain largely unclear.

In human cancers, ALT activation is intimately linked to the mutational status of the chromatin modulator genes *ATRX* and *DAXX* ^9-13^. Functionally, ATRX and DAXX are known to form a histone H3.3-specific chaperone complex that facilitates replication-independent nucleosome assembly at heterochromatic regions including telomeres ^14-18^. A survey of ~ 7,000 patient samples in 31 cancer types found that 5% of them harbor genetic alterations of *ATRX* or *DAXX* that also concurrently present ALT features ^19^. This tight correlation has raised the possibility that ALT activation is a consequence of histone management dysfunction.

The ALT mechanism relies on homologous recombination-directed telomere DNA synthesis. Cumulative evidences suggest that ALT activation emanates from telomere replication stress and stalled replication forks ^5, 20, 21^. Indeed, depletion of ATRX or DAXX disrupts replication-independent nucleosome incorporation and induces telomere chromatin de-condensation that progressively activate the homology-directed telomere repair pathway ^22^. As a consequence, the homologous repair-based ALT mechanism becomes the only viable path for the *ATRX* or *DAXX* mutant cells to achieve replicative immortality. In this sense, *ATRX* or *DAXX* loss, while promoting tumorigenesis through activating the ALT-directed telomere maintenance pathway, also simultaneously creates an intrinsic telomere replication defect that can potentially be explored for synthetic lethal-like interactions.

Through CRISPR/Cas9-based genetic screens of paired isogenic ALT and TERT-immortalized control cells lines, here we identify histone demethylase KDM2A as a selective molecular vulnerability of cells that depend on ALT-directed telomere maintenance. We demonstrate that KDM2A functions to facilitate dissolution of the ALT-specific multitelomere clusters following homology-based telomere synthesis. We further show that KDM2A promotes ALT multitelomere de-clustering by facilitating SUMO isopeptidase SENP6-mediated SUMO deconjugation at telomeres. These findings together establish KDM2A as a promising therapeutic target for ALT-dependent cancers.

## Results

### A CRISPR-based genetic screen of chromatin regulators required by ALT cells

To uncover the molecular vulnerabilities of cells that rely on alternative lengthening of telomere maintenance, we developed isogenic pairs of ALT cell lines from ATRX-depleted human lung IMR90 fibroblasts following our established immortalization protocol (Fig. 1a) ^22^. Compared to the control IMR90-T cell lines that were immortalized by telomerase (TERT) expression, these ALT-immortalized IMR90 cells exhibited highly elevated levels of 53BP1-associated telomere dysfunction-induced foci (TIF) (Supplementary Fig. 1a, b), consistent with the notion that ALT telomeres experience chronic replication stress and are intrinsically unstable ^5, 12, 23, 24^.

**Fig. 1.**
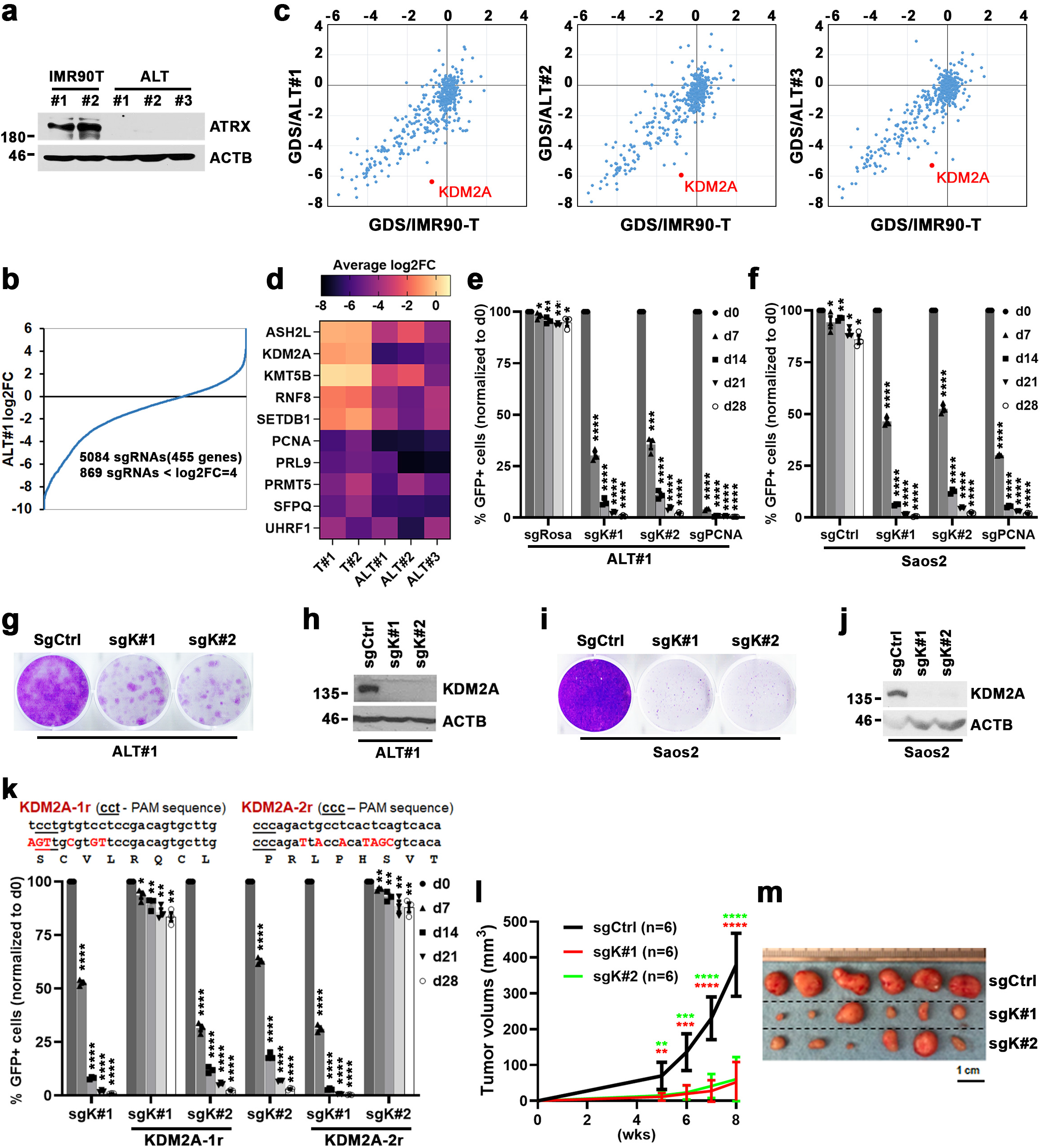
CRISPR-based screens identify KDM2A as selectively essential in ALT-dependent cells. **a** Western blot analysis of ATRX protein expression in whole-cell lysates prepared from the indicated cells. **b** Ranking of sgRNAs by log2 fold-change (log2FC) of abundance (ratio of start to end point) in ALT#1 cells. x axis shows targeting sgRNAs; y axis shows the log2FC of each targeting sgRNA after 16 population doublings. **c** Gene dependency scores (GDS) in IMR90-T (x axis) versus ALT#1, ALT#2, or ALT#3 cells (y axis). The GDS was calculated by averaging the log2FC of all sgRNAs targeting that gene. **d** Heatmap depicts log2FC of average sgRNA abundance of selected genes in indicated cells after 16 population doublings. **e, f** Competition-based proliferation assay of *KDM2A*-targeted sgK#1 and sgK#2 in ALT#1 (**e**) or Cas9-expressing Saos2 cells (**f**). A GFP reporter is linked to sgRNA expression. Plotted is the %GFP cells (normalized to the d0 measurement) at the indicated time-points. The non-targeting sgCtrl was included as a negative and sgRNA targeting *PCNA* as a positive control. **g, h** Clonogenic assay (**g**) and KDM2A western blot analysis (**h**) of sgCtrl, sgK#1 or sgK#2-transduced ALT#1 cells. Crystal violet staining was conducted at day 20 post seeding. **i, j** Crystal violet-based clonogenic survival assay (**i**) and KDM2A western blot analysis (**j**) of sgCtrl, sgK#1 or sgK#2-transduced Saos2 cells. Crystal violet staining was conducted at day 24 post seeding. **k** Competition-based proliferation assay of GFP-linked sgRNA in Saos2 cells complemented with either empty vector control or the CRISPR-resistant *KDM2A* synonymous mutants (*KDM2A-1r* or *KDM2A-2r*). Note, the *KDM2A-1r*-transduced cells are resistant to sgK#1 but sensitive to sgK#2 action; the *KDM2A-2r*-transduced cells are resistant to sgK#2 but sensitive to sgK#1 action. **l** Tumor growth curves of sgCtrl, sgK#1 or sgK#2-transduced Saos2 cells. Data are expressed as means ± s.e.m. of 6 biological replicates, unpaired t-test. **m** Image of tumors collected at week 8 post subcutaneous transplantation. In (**e**), (**f**) and (**k**), Data are expressed as means ± s.e.m. of three independent experiments. * denotes ns; ** P < 0.05; *** P < 0.01; **** P < 0.001; paired t-test.

To profile chromatin regulators that are selectively required for ALT-immortalized cells, we constructed a library that contained ~5,000 sgRNAs targeting 455 chromatin modifiers, readers, and effectors (~10 sgRNAs/gene) as well as ~ 100 control sgRNAs. To enhance the targeting efficiency, the sgRNAs were designed using an algorithm linked to protein domain annotation. The genetic screens were conducted by transducing the sgRNA library into three independently derived ALT-immortalized IMR90 cells (hereafter referred to as ALT#1, ALT#2 and ALT#3) and two paired isogenic control IMR90-T cells that were immortalized by TERT expression (referred to as IMR90-T#1 and #2). The pools of library-transduced cells were passaged for 16 population doublings before being subjected to next generation sequencing-based quantification. The relative effect of each sgRNA on cell growth was scored by calculating the log_2_ fold change (log2FC) of sgRNA abundances at the beginning and end of the culture periods (Fig. 1b; Supplementary Fig. 2a). The spike-in positive (sgPCNA, sgRPA3, sgCDK1, sgCDK9 sgTIP60, and sgTTF2) and non-targeting negative (sgNeg1 - 100) control sgRNAs served as quality controls to validate the overall accuracy of the screening strategy.

For each gene, we calculated its gene dependency score (GDS) by averaging the log2FC of all targeting sgRNAs (4 - 14 per gene) (Supplementary Table 1). To rank the priority of ALT-selective vulnerabilities, GDS of individual genes in indicated ALT cell lines (x-axis) were plotted against their scores in control IMR90-T cells (y-axis) (Fig. 1c). Unsurprisingly, many of the identified gene dependencies were pan-essential and scored comparably in the ALT and their control IMR90-T cells. Among the genes selectively required for ALT cells were several genes with diversified chromatin regulatory functions, including *ASM2L, KDM2A, KMT5B, RNF8*, and *SETDB1* (Fig. 1d). The most prominent hit in this screen was *KDM2A*, a member of the Jumonji C (Jmjc) domain-containing histone lysine demethylase family that targets lower methylation states of H3K36 (Kme1 and Kme2) but has no known function in telomeres ^25, 26^. Notably, the majority (7 of 12) of sgRNAs targeting *KDM2A* in the library screens caused robust growth inhibition phenotypes in all three ALT lines (log2FC < -5.0) but were relatively ineffective against control IMR90-T lines (Supplementary Fig. 2b), suggesting that KDM2A is selectively essential for ALT cells.

To validate the pooled screens, we next analyzed the proliferative impact of individual sgRNAs through fluorescence-activated cell sorting (FACS)-based competition assays ^22^. Consistent with the screen result, targeting *KDM2A* using two newly designed sgRNAs (referred to as sgK#1 and sg#2) profoundly suppressed growth of the ALT-dependent ALT#1 (Fig. 1e), osteosarcoma Saos2 (Fig. 1f), U2OS (Supplementary Fig. 2c), G292 (Supplementary Fig. 2d), rhabdomyosarcoma Hs729 (Supplementary Fig. 2e), and patient-derived pGBM6 glioblastoma cells (Supplementary Fig. 2f). As a complementary approach, we conducted crystal violet-based clonogenic growth assays. Consistently, sgK#1 or sgK#2-mediated KDM2A ablation in ALT#1 (Fig. 1g, h), Saos2 (Fig. 1i, j), or U2OS cells (Supplementary Fig. 2g, h) greatly inhibited their clonogenic growth. Notably, the competition- and clonogenic-based proliferation assays demonstrated that complementation of the sgK#1- or sgK#2-resistant *KDM2A* cDNAs fully rescued the growth inhibition caused by the respective sgRNAs (Fig. 1k; Supplementary Fig. 2i-l), confirming their on-target effect.

Finally, to assess whether KDM2A is essential for ALT cell growth in vivo, Saos2 cells transduced with sgCtrl or *KDM2A*-specific sgRNAs were subcutaneously grafted into immunocompromised recipient mice. Analysis of tumor growth revealed that inactivation of KDM2A by sgK#1 or sgK#2 significantly inhibited the in vivo tumorigenicity (Fig. 1l, m), indicating that KDM2A is required for ALT-dependent tumor propagation.

### KDM2A is not essential for non-ALT cell proliferation and survival

To ascertain whether KDM2A is selectively essential for ALT cells, we next examined the growth effect of KDM2A depletion on the wild-type *ATRX* cDNA-complemented U2OS cells. In line with the previous study ^27^, re-expression of ATRX in the ATRX-null U2OS cells suppressed ALT and APB formation (Supplementary Fig. 3a-c). Moreover, targeting the ATRX-complemented U2OS with sgK#1 or sgK#2 caused a much attenuated growth inhibition phenotype as compared to the parental U2OS cells (Supplementary Fig. 3d), indicating that KDM2A is selectively essential for ALT cells. Consistently, ectopic expression of wild-type *DAXX* cDNA in the *DAXX*-mutant G292 cells suppressed APB formation and also rescued the growth inhibition caused by *KDM2A*-specific sgRNAs (Supplementary Fig. 4e-h) ^28, 29^.

To further assess the selectivity of KDM2A dependencies, we next conducted competition-based proliferation assays in a panel of non-ALT cells of diversified tissue origins. Compared to ALT-dependent cells in which ablation of KDM2A by sgK#1 or #2 led to 35 - 100 folds drop-out in a period of 4 weeks, targeting *KDM2A* by the same sgRNAs incurred noticeable but minor growth inhibition effects (< 2.5 folds drop-out) in the panel of non-ALT human cell lines, including IMR90-T (Fig. 2a), HeLa of cervical cancer (Fig. 2b), NCI-H1299 of non-small cell lung carcinoma (Fig. 2c), MG63 of osteosarcoma (Supplementary Fig. 4a), MCF7 of breast cancer (Supplementary Fig. 4b), glioma cell lines A172, LN464 and U118 (Supplementary Fig. 4c-e), and primary lung fibroblast IMR90 cells (Supplementary Fig. 4f). As the positive control, sgRNAs targeting *PCNA* induced growth arrest in all tested cell lines, ruling out the possibility of inefficient genome editing. These results were further verified by clonogenic assays of sgK#1 or #2-transduced IMR90-T (Fig. 2d, e), HeLa (Fig. 2f, g), or NCI-H1299 cells (Fig. 2h, i).

**Fig. 2.**
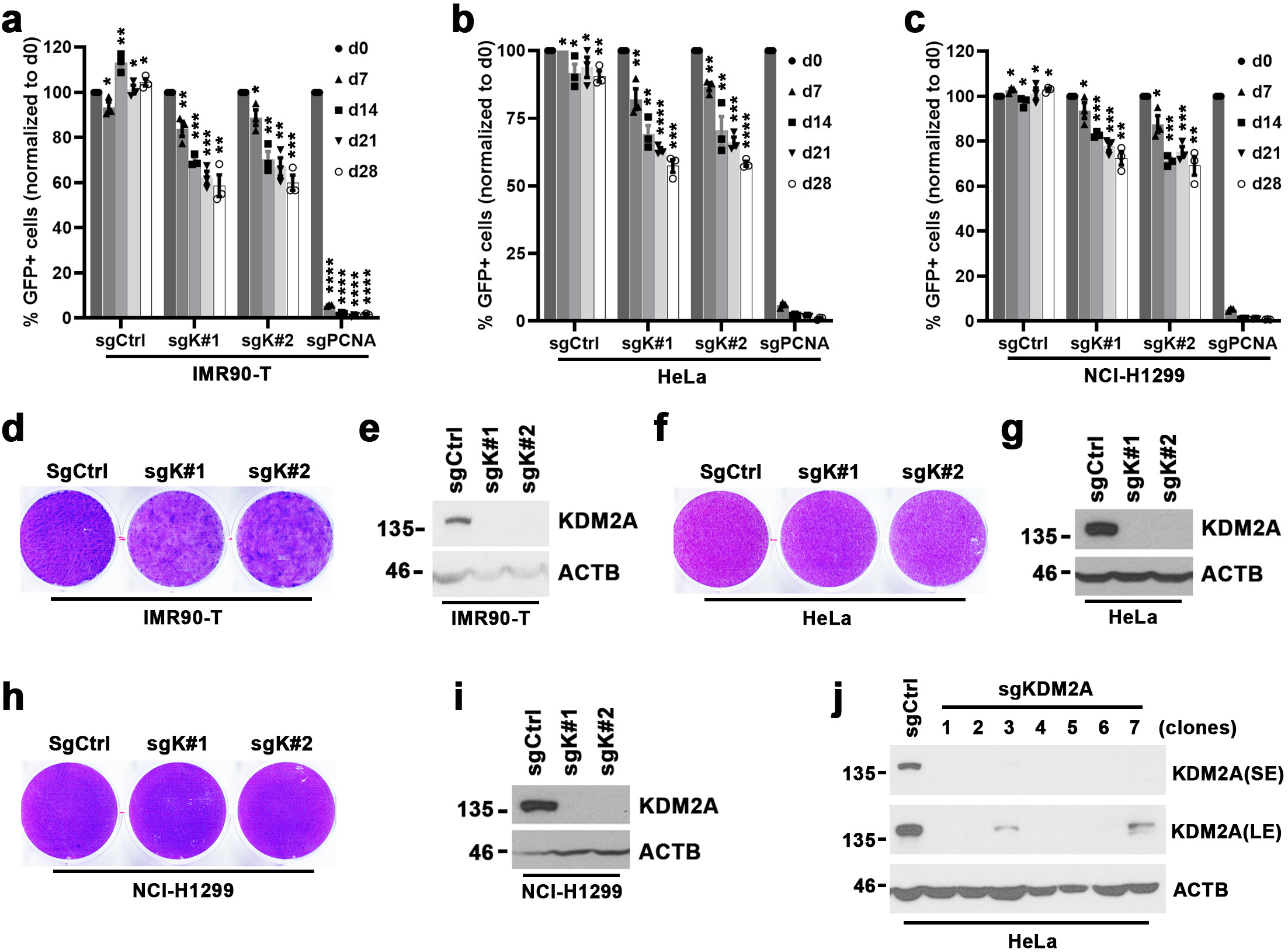
KDM2A is dispensable for non-ALT cells. **a-c** Competition-based proliferation assays of the indicated sgRNAs in IMR90-T (**a**), HeLa (**b**), or NCI-H1299 cells (**c**). A GFP reporter is linked to sgRNA expression. sgCtrl and sgPCNA were included as a negative or a positive control respectively. Data are expressed as means ± s.e.m. of three independent experiments. * denotes ns; ** P < 0.05; *** P < 0.01; **** P < 0.001; paired t-test. **d, e** Clonogenic assay (**d**) and KDM2A western blot analysis (**e**) of sgCtrl, sgK#1 or sgK#2-transduced IMR90-T cells. Crystal violet staining was conducted at day 22 post seeding. **f, g** Clonogenic assay (**f**) and KDM2A western blot analysis (**g**) of sgCtrl, sgK#1 or sgK#2-transduced HeLa cells. Crystal violet staining was conducted at day 15 post seeding. **h, i** Clonogenic assay (**h**) and KDM2A western blot analysis (**i**) of sgCtrl, sgK#1 or sgK#2-transduced NCI-H1299 cells. Crystal violet staining was conducted at day 15 post seeding. **j** Western blot analysis of KDM2A and ACTB in whole-cell lysates prepared from indicated HeLa cell lines. The KDM2A depleted lines (clone#1-7) were established from clonally isolated HeLa cells transduced with sgK#1. The sgCtrl transduced HeLa line was included as a control. SE, short exposure; LE, long exposure.

To determine whether KDM2A protein expression is dispensable for non-ALT cell survival, we next applied the CRISPR/Cas9 system to delete KDM2A in HeLa and LN464 cells. By western blot survey of the panels of clonally derived cultures, we identified 5 (out of 17) HeLa and 4 (out of 15) LN464 clones that were completely depleted of KDM2A protein expression (Fig. 2j; Supplementary Fig. 4g). These KDM2A-null cells were viable and proliferated at slightly slower rates than the parental HeLa or LN464 cells (Supplementary Fig. 4h, i), indicating that *KDM2A* is not a pan-essential gene. By contrast, we were not able to recover any surviving KDM2A-null clones from sgK#1- or sgK#2-transduced ALT#1, Hs729, Saos2, or U2OS cell cultures. These findings support KDM2A as a potential therapeutic target selectively for ALT-dependent human cancers.

### ALT cell growth depends on multiple functional activities of KDM2A

As a modular protein, KDM2A consists of a variety of structural motifs that serve different activities (Fig. 3a). To map the KDM2A protein domains critical for its ALT-supporting function, we synthesized a CRISPR exon-tilling library that comprised 492 sgRNAs targeting the entire *KDM2A* open reading frame (Supplementary Table 2). The tiling library was transduced into Saos2, ALT#1, ALT#2, and control IMR90-T cells. Using massively parallel sequencing, we calculated the depletion fold of each sgRNA over 16 population doublings. Among the sgRNAs that induced the most robust growth inhibition phenotypes in ALT-dependent Saos2, ALT#1, and ALT#2, were enriched for ones that target the N-terminal demethylase domain, CXXC-type zinc finger (ZnF), or PHD domain of KDM2A (Fig. 3b, c; Supplementary Fig. 5a), indicating that DNA binding and demethylase activities of KDM2A are required for mediating its ALT cell supporting function. By contrast, the sgRNAs that target the C-terminal domain, F-Box and other linker regions had less pronounced growth inhibition effects, suggesting that the E3 ligase activity of KDM2A is likely not required for sustaining ALT cell growth. Finally, none of the sgRNAs induced robust dropout phenotypes in the control IMR90-T cells (Supplementary Fig. 5b), further supporting that KDM2A is a selective vulnerability of ALT-dependent cells.

**Fig. 3.**
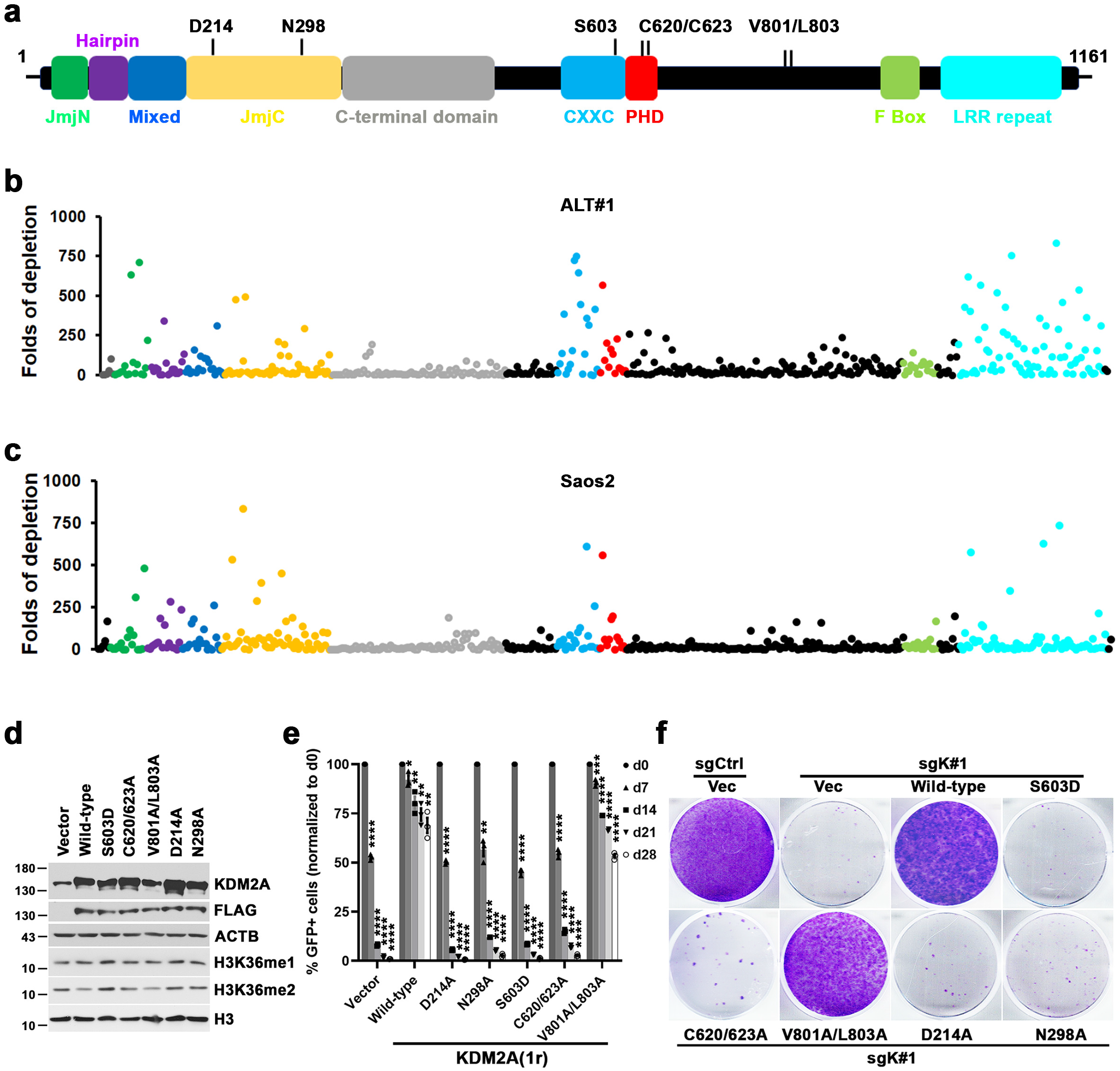
DNA binding and demethylase activities of KDM2A are both required for its ALT supporting function. **a** Schematic of full-length KDM2A protein. The domains are labeled and color-coded. **b, c** CRISPR-based KDM2A tiling assay in ALT#1 (**b**) and Saos2 cells (**c**). Plotted is the fold changes of sgRNA abundance (ratio of start to end point) after 16 population doublings in culture. x axis shows targeting sgRNAs and the domain location of each sgRNA within KDM2A protein is indicated by colors; y axis shows the fold changes of each targeting sgRNA following the culture period. **d** Western blot analysis of KDM2A, Flag and ACTB in whole-cell lysates prepared from Saos2 cells transduced with indicated constructs. The Flag-tagged wild-type and KDM2A mutants were generated from CRISPR-resistant *KDM2A-1r* construct. **e** Competition-based proliferation assay of *KDM2A*-targeted sgK#1 (linked with GFP expression) in Saos2 cells transduced with the indicated constructs. The bar graphs are expressed as means ± s.e.m. of three independent experiments, paired t-test. * denotes ns; ** P < 0.05; *** P < 0.01; **** P < 0.001. **f** Clonogenic assay of sgK#1 in Saos2 cells transduced with the indicated CRISPR-resistant wild-type or *KDM2A* mutants. Crystal violet staining was conducted at day 24 post cell seeding.

To validate the exon-tiling scan results, we next transduced Sao2 cells with sgK#1-resistant *KDM2A* cDNAs encoding wild-type or mutants defective of DNA binding (S603D) ^30, 31^, PHD domain structural integrity (C620/623A) ^30^, HP1 protein interaction (V801A/L803A) ^30^, or demethylase activity (D214A or N298A) ^25^. Noticeably, despite being expressed at comparable levels, western blot analysis found that only cells transduced with wild-type or the V801A/L803A mutant showed visible reduction of H3K36me2 as compared to the vector transduced control cells (Fig. 3d). Consistently, competition and clonogenic-based proliferation assays revealed that complementation of wild-type or V801A/L803A, but not other mutants, fully rescued sgK#1-induced growth inhibition in Saos2 cells (Fig. 3e, f), indicating that HP1 protein interaction activity is dispensable for its ALT supporting function.

The other identified KDM2A essential region in the exon-tilling scan of ALT cells was the C-terminal leucine-rich repeats (LRR). To test its function in ALT cell growth, we constructed a KDM2A ΔLRR mutant (aa 1-aa 945) that is deleted of LRR motif. Consistent with the exon-tiling scan results, a competition-based proliferation assay indicated that complementation of the CRISPR-resistant ΔLRR mutant was not able to rescue the sgK#1-induced growth inhibition phenotype in Saos2 cells (Supplementary Fig. 6a, b).

### KDM2A binds physically to ALT telomeres

In human cancers and immortalized cell lines, ALT activation is closely associated with genetic alterations that affect the histone H3.3 chaperone ATRX-DAXX complex ^11, 12, 19, 32^. Indeed, a western blot survey of cell lines used in this study revealed that ATRX protein was broadly expressed in the non-ALT cells but absent in ALT-dependent cells (Fig. 4a). By contrast, KDM2A protein was expressed across the panel of cell lines regardless of their tissues of origin or ATRX expression status, suggesting that their selective KDM2A dependency is not due to differential protein expression.

**Fig. 4.**
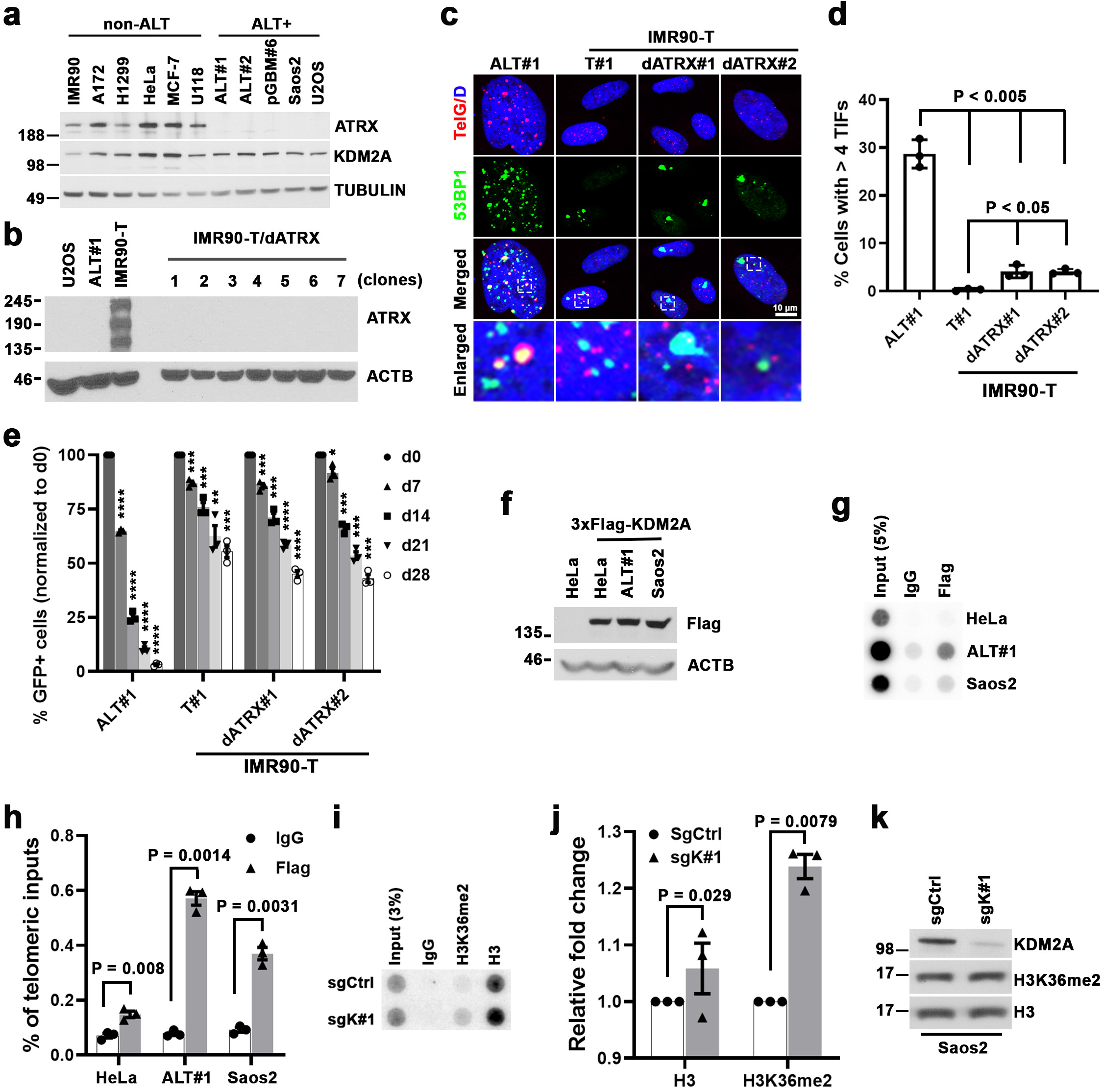
KDM2A regulates H3K36me2 at ALT telomeres. **a** Western blot analysis of ATRX protein expression in whole-cell lysates prepared from the indicated non-ALT and ALT-dependent (ALT+) cells. **b** Western blot analysis of ATRX expression in whole-cell lysates prepared from control or ATRX-depleted IMR90-T cells. The ATRX depleted lines (dATRX clone#1-7) were established from clonally isolated IMR90-T cells transduced with ATRX-targeted sgRNA. The lysates from U2OS and ALT#1 cells were included as negative controls of ATRX expression. **c** Representative immuno-FISH images of 53BP1 and telomeres in ALT#1, IMR90-T (T#1), or ATRX-depleted IMR90-T (dATRX#1 and dATRX#2) cells. Scale bar, 10 μm. **d** Percentages of cells containing ≥ 4 53BP1-associated telomere dysfunction-induced foci (TIFs). Data are expressed as means ± s.e.m. of three independent experiments, unpaired t-test. **e** Competition-based proliferation assay of KDM2A-targeted sgK#1 (linked with GFP expression) in the indicated cell lines. Data are expressed as means ± s.e.m. of three independent experiments. **f** Western blot analysis of Flag and ACTB (loading control) using whole-cell lysates prepared from control or Flag-tagged wild-type KDM2A-transduced HeLa, ALT#1 or Saos2 cells. **g, h** Telomere dot blot analysis (**g**) and quantification (**h**) of anti-Flag or IgG chromatin immunoprecipitation (ChIP) in the indicated cell lines. The IgG ChIP was included as a control for non-specific signal. The input and ChIP DNAs processed against the indicated antibodies were assayed by dot-blotting and hybridized with ^32^P-labeled TelG probe. The relative enrichment was calculated after normalization of ChIP DNA signals to the respective input DNA signals. Data are expressed as means ± s.e.m. of three independent experiments, paired t-test. **i, j** Telomere dot blot analysis (**i**) and quantification (**j**) of anti-H3K36me2, anti-H3 or IgG ChIP in sgCtrl or sgK#1-transduced Saos2 cells. The relative enrichment was calculated after normalization to the respective ChIP DNA signals of sgCtrl-transduced samples. Data are expressed as means ± s.e.m. of three independent experiments, paired t-test. **k** Western blot analysis of KDM2A, H3K36me2, and H3 in cell lysates prepared from sgCtrl or sgK#1-transduced Saos2 cells.

To explore whether KDM2A is a synthetic vulnerability of ATRX deficiency independently of its ALT status, we generated the *ATRX* knockout cells from TERT-transduced IMR90-T or LN464-T cells (Fig. 4b; Supplementary Fig. 7a). These clonally derived ATRX-null IMR90-T or LN464-T cells (referred to as dATRX#1 and #2) were viable and proliferated at slightly slower rates than their respective control cells (Supplementary Fig. 7b, c). Consistent with our previous findings ^22^, these TERT-overexpressed ATRX-null cells displayed low levels of telomere dysfunction and TIF formation (Fig. 4c, d; Supplementary Fig. 7d, e). Competition-based proliferation assays further revealed that targeting these ATRX-knockout IMR90-T or LN464-T cells by *KDM2A-*specific sgRNAs only moderately affected their fitness, which is in stark contrast to ALT#1 cells where KDM2A inhibition caused robust growth arrest (Fig. 4e; Supplementary Fig. 7f). These findings suggest that KDM2A is not simply a synthetic lethal vulnerability of ATRX deficiency.

Notably, among the 21 histone lysine demethylases reported in the literature, our screen identified KDM2A as the only one essential for ALT cell growth (Supplementary Fig. 7g). To examine whether KDM2A may act physically at telomeres, we transduced Flag-tagged KDM2A into Saos2, ALT#1 and ALT-null HeLa cells. Consistent with a previous study ^33^, telomere dot-blot analysis of anti-Flag chromatin immunoprecipitation (ChIP) of the transduced cells revealed significant enrichment of KDM2A at telomeres, preferentially in ALT cells (Fig. 4f-h). By comparison, anti-Flag ChIP/Alu dot-blot analysis of the chromatin immunoprecipitation did not detect ALT-preferential enrichment (Supplementary Fig. 8a, b). Finally, anti-H3K36me2 ChIP/telomere dot-blot analysis of KDM2A-depleted Saos2 and ALT#1 cells revealed significantly increased levels of telomere H3K36me2 as compared to the sgCtrl-transduced cells (Fig. 4i-k; Supplementary Fig. 8c-e). These results collectively support a direct involvement of KDM2A in ALT telomere maintenance.

### KDM2A facilitates chromosomal segregation of ALT cells

ALT cells are characterized by persistent telomere DNA replication stress and rely on homology-based DNA repair pathways to elongate their telomeres ^5, 8, 23^. To interrogate how KDM2A depletion affects ALT cell growth, we conducted cell cycle analysis and found a markedly elevated G2/M phase accumulation in KDM2A-depleted ALT#1 or Saos2 cells as compared to their respective control cells (Fig. 5a; Supplementary Fig. 9a, b). But despite its selective role in growth regulation of ALT cells, depletion of KDM2A did not seem to significantly affect many ALT-associated activities. For example, quantitation of control and KDM2A-depleted ALT#1 or Saos2 cells showed comparable levels of APB formation (Fig. 5b, c; Supplementary Fig. 9c, d), a hallmark of ALT activation ^34^. Similarly, analysis of telomere length and C-rich extrachromosomal telomeric repeats (C-circles) in control and KDM2A-depleted ALT#1 or Saos2 cells also revealed no significant changes in telomere length heterogeneity and C-circle formation (Fig. 5d, e; Supplementary Fig. 9e, f).

**Fig. 5.**
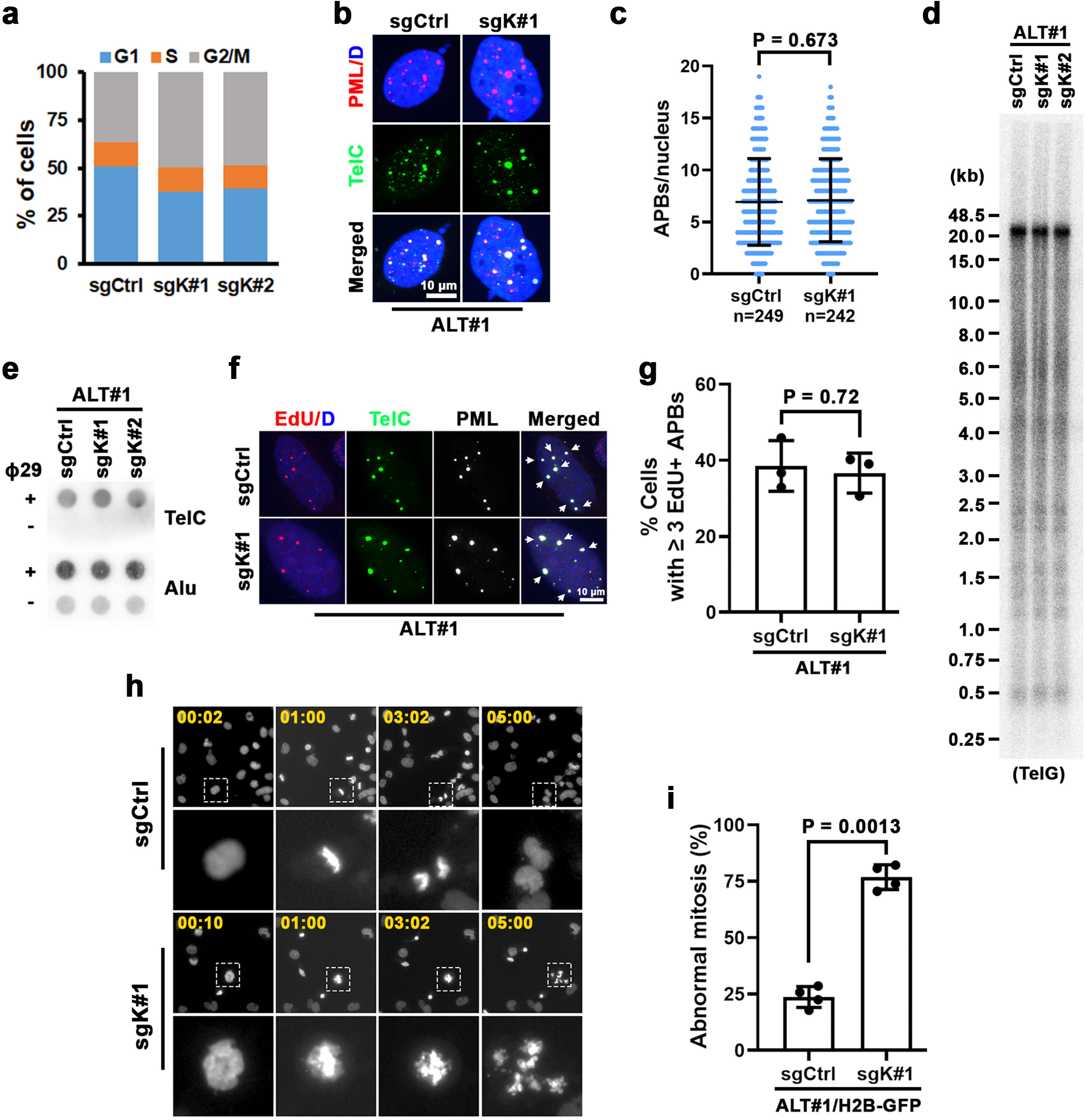
KDM2A depletion disrupts ALT chromosomal segregation and mitotic division. **a** Cell cycle distribution analysis of sgCtrl, sgK#1 or sgK#2-transduced ALT#1 cells. **b** Representative immuno-FISH images of ALT-associated PML bodies (APBs) in sgCtrl or sgK#1-transduced ALT#1 cells. Scale bar = 10 µm. **c** Percentages of cells containing ≥ 4 APBs are expressed as means ± s.e.m. of three independent experiments, unpaired t-test. **d** Telomere restriction fragment analysis of telomere length of sgCtrl, sgK#1 or sgK#2-transduced ALT#1 cells. Genomic DNAs prepared from the indicated cells were assayed by ^32^P-labeled TelG probe. **e** C-circle assays of sgCtrl, sgK#1 or sgK#2-transduced ALT#1 cells. Genomic DNAs were prepared from the indicated cells and assayed by ^32^P-labeled TelC probe. **f** Representative immuno-FISH images of EdU colocalized APBs (EdU-APBs) in sgCtrl or sgK#1-transduced ALT#1 cells. Cells were synchronized in G2 with sequential thymidine and CDK1 inhibitor treatment and then labeled with EdU for 2 h. EdU was assayed by Click-It reaction, PML was analyzed by IF, and telomeres were detected by FISH. The arrows denote EdU-APB foci. **g** Percentages of cells containing ≥ 3 EdU-APB foci. Data are expressed as means ± s.e.m. of three independent experiments, unpaired t-test. **h** Representative frames of time-lapse fluorescence live cell imaging of GFP-H2B-expressing ALT#1 cells transduced with sgCtrl or sgK#1. The cells were synchronized in G2 with sequential thymidine and CDK1 inhibitor treatment before timed release into M phase. Time in minutes is shown in upper left corners. **i** Percentages of aberrant mitosis are expressed as means ± s.e.m. of four independent experiments, unpaired t-test.

To investigate whether ablation of KDM2A affects ALT telomere DNA synthesis, sgCtrl and sgK#1-transduced ALT#1 cells were synchronized to G2 phase with sequential thymidine and CDK1 inhibitor Ro-3306 treatment before 5-ethynyl-2’-deoxyuridine (EdU) labeling. Interestingly, the following assay of ALT telomere DNA synthesis in APBs (ATSA) found no significant differences in their levels of telomere EdU incorporation (Fig. 5f, g). A similar observation was made in comparing sgCtrl and sgK#1-transduced Saos2 cells (Supplementary Fig. 9g, h), suggesting that KDM2A may act downstream of homology-directed telomere DNA synthesis.

KDM2A depletion in ALT cells induces G2/M phase accumulation. This abnormal cell cycle distribution could be caused by cell cycle arrest or dysfunctional mitosis.. To monitor mitotic cell division by time-lapse live cell imaging, the GFP-H2B-expressing ALT#1 cells transduced with sgCtrl or sgK#1 were synchronized by sequential thymidine and CDK1 inhibitor treatment before timed release. For the control sgRNA-transduced ALT#1 cells, we found that a majority (102/134; 76%) of them that had entered mitosis during the imaging periods underwent normal chromosomal segregation and cytokinesis (Supplementary Movie 1). By comparison, of the 112 KDM2A-depleted cells that were tracked for their mitotic division, 74% (83 of 112) of them displayed aberrant chromosomal segregation and eventually submitted to mitotic catastrophe, as indicated by their hyper-condensed chromatin aggregates and/or DNA fragmentation (Fig. 5h, i; Supplementary Movie 2). These gross mitotic failures were also visualized in the live imaging of KDM2A depleted ALT#2 cells that stably expressed GFP-H2B (Supplementary Fig. 10a, b; Supplementary Movies 3, 4). By contrast, depletion of KDM2A in GFP-H2B-expressing IMR90-T cells minimally affected their mitotic division (Supplementary Fig. 10c, d; Supplementary Movies 5, 6).

Mitotic catastrophe represents a regulated mechanism that responds to aberrant mitoses by removing damaged cells from the cycling population ^35, 36^. Indeed, analysis of mitotic outcomes of the KDM2A-depleted ALT#1 cells indicated a significantly elevated mitotic death (Supplementary Fig. 10e, f). Western blot analysis further revealed an increased level of apoptosis in sgK#1-transduced ALT#1 cells, as evidenced by their cleaved PARP1 production (Supplementary Fig. 10g). These findings suggest that KDM2A functions to facilitate chromosomal segregation of ALT cells and its loss induces apoptosis-dependent mitotic death.

### KDM2A is required for ALT multitelomere chromatin de-clustering

ALT-directed telomere synthesis occurs within APBs where recombinogenic telomeres from different chromosomes coalesce together ^8, 37, 38^. This process is cell cycle-regulated, and the clustered telomeres must be disassembled prior to anaphase to ensure proper chromosome segregation ^38^. To test whether KDM2A might be involved in the regulation of ALT telomere de-clustering following recombination-directed synthesis, we synchronized the sgCtrl and sgK#1-transduced ALT#1 cells to G2 phase. The immuno-FISH analysis of the G2-synchronized cells revealed comparable levels of APB formation (Fig. 6a-c), indicating that KDM2A loss did not significantly affect telomere clustering and APB assembly. Following timed release from the CDK1 inhibitor block, the majority of control ALT#1 cells that entered mitosis and were marked by H3-Ser10 phosphorylation (pH3S10) had cleared their multitelomere clusters (Fig. 6a-c). By comparison, ~ 73% of mitotic ALT#1 cells depleted of KDM2A retained clustered telomere foci, indicating a defect in telomere declustering. And those faulty chromosomes eventually underwent aberrant segregation and mitotic catastrophe (Fig. 6d). Similar phenotypes were also observed in KDM2A-depleted Saos2 (Supplementary Fig. 11a-d), ALT#2 (Supplementary Fig. 11e-g), and G292 cells (Supplementary Fig. 11h-j).

**Fig. 6.**
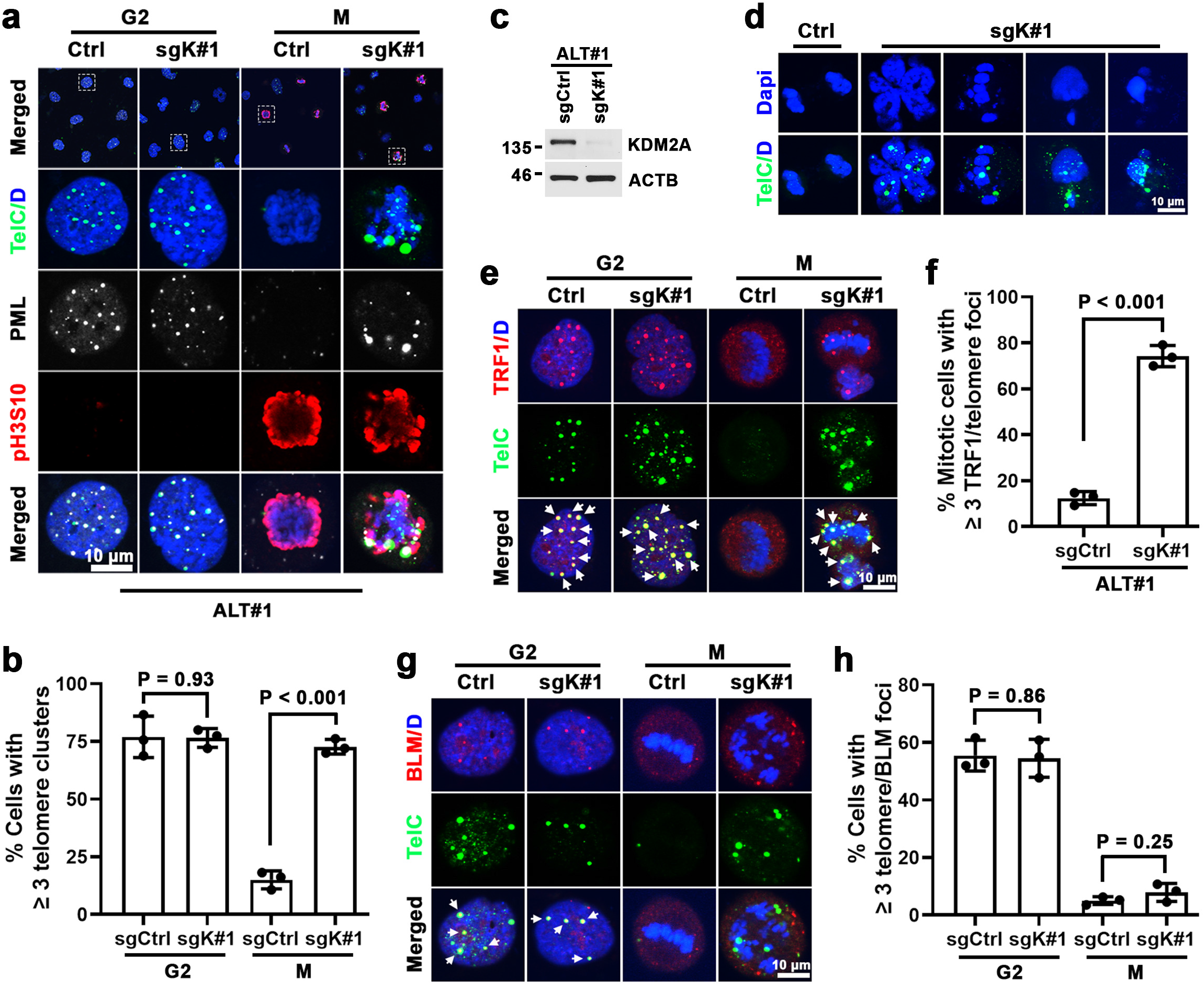
KDM2A promotes ALT telomere de-clustering after recombination. **a** Representative immuno-FISH images of multitelomere clusters in G2 or mitotic ALT#1 cells transduced with sgCtrl or sgK#1. The G2 cells were synchronized from sequential thymidine and CDK1 inhibitor treatment. The mitotic (M) cells were from G2 synchronized cells upon 1 h release from CDK1 inhibitor. PML and mitotic marker phospho-Histone H3-Ser10 (pH3S10) were analyzed by IF and telomeres were detected by FISH. **b** Percentages of cells containing ≥ 3 multitelomere cluster foci. **c** Western blot analysis of KDM2A and ACTB in whole-cell lysates prepared from sgCtrl or sgK#1-transduced ALT#1 cells. **d** Representative images of abnormal mitotic multitelomere clusters in sgK#1-transduced ALT#1 cells. G2 synchronized ALT#1 cells transduced with sgCtrl or sgK#1 were released into mitosis for 2 h. Telomeres were detected by FISH. **e** Representative immuno-FISH images of TRF1-telomere association in G2 or M-phase ALT#1 cells transduced with sgCtrl or sgK#1. The M-phase cells were from G2 synchronized cells upon 1 h release from CDK1 inhibitor. TRF1 was analyzed by IF and telomeres were detected by FISH. The arrows denote TRF1-associated telomere foci. **f** Percentages of mitotic cells containing ≥ 3 TRF1-associated telomere foci. **g** Immuno-FISH analysis of BLM and telomeres colocalization in G2- or M-phase ALT#1 cells transduced with sgCtrl or sgK#1. BLM was analyzed by IF and telomeres were detected by FISH. The arrows denote BLM-associated telomere foci. **h** Percentages of cells containing ≥ 3 BLM-associated telomere foci. Note, all bar graphs are expressed as means ± s.e.m. of three independent experiments, unpaired t-test. Scale bar, 10 μm.

To further validate KDM2A’s role in ALT telomere declustering, we conducted a reconstitution experiment. As expected, complementation of CRISPR-resistant *KDM2A* cDNAs into ALT#1 cells fully rescued the sgK#1-induced telomere segregation defects (Supplementary Fig. 11k-m). Moreover, re-expression of ATRX in ALT#1 cells suppressed KDM2A depletion-induced multitelomere cluster formation and segregation dysfunction (Supplementary Fig. 12a-c), confirming that KDM2A is required for post-recombinational ALT telomere declustering.

To characterize those aberrantly retained M-phase multitelomere clusters, we next analyzed their association with telomere binding proteins. In line with the previous reports ^38, 39^, telomeres in the control mitotic ALT#1 cells were largely condensed and exhibited reduced interaction with telomere binding protein TRF1 (Fig. 6e, f). By contrast, the aberrantly clustered mitotic telomeres in the KDM2A depleted ALT#1 cells were still strongly associated with TRF1, suggesting compromised condensation at these telomeres. Similar findings were also obtained in KDM2A-depleted Saos2 cells (Supplementary Fig. 13a, b).

Intermediates of homologous recombination are potential sources of chromosome mis-segregation if not removed before anaphase ^40, 41^. Since ALT functions through homologous recombination-based telomere DNA synthesis, we next asked whether the aberrant multitelomere chromatin clusters were unresolved recombination intermediates. The Bloom’s syndrome protein BLM is a RecQ family helicase that drives ALT-associated telomere synthesis and intermediate telomere structure processing ^41-44^. Notably, immuno-FISH analysis of G2-synchronized control and KDM2A-depleted ALT#1 or Saos2 cells revealed comparable levels of telomere-associated BLM foci formation (Fig. 6g, h; Supplementary Fig. 13c, d). Upon mitotic entry following release from the CDK1 inhibitor block, telomeres in the sgCtrl-transduced control cells as expected was dissociated of BLM binding. Similarly, BLM was also completely cleared from the aberrant telomere clusters of mitotic ALT#1 or Saos2 cells depleted of KDM2A, indicating that these abnormal telomere clusters are unlikely to be unresolved DNA recombination intermediates.

### KDM2A promotes SENP6-mediated ALT telomere de-SUMOylation

SMC5/6 complex-activated protein SUMOylation is critical for ALT-directed multitelomere clustering and APB formation ^45^. Consistently, we found that SMC5 co-localized with telomeres at the PML bodies of ALT#1 cells and its knock-down blocked telomere recruitment to PML bodies and ALT telomere cluster foci formation (Supplementary Fig. 14a, b). Moreover, compared to scrambled control siRNA treatment, transduction of *SMC5* siRNA in sgK#1-transduced ALT#1 cells strongly attenuated the KDM2A depletion-induced aberrant M-phased telomere cluster formation (Fig. 7a-c), indicating that KDM2A functions downstream of SMC5/6 action.

**Fig. 7.**
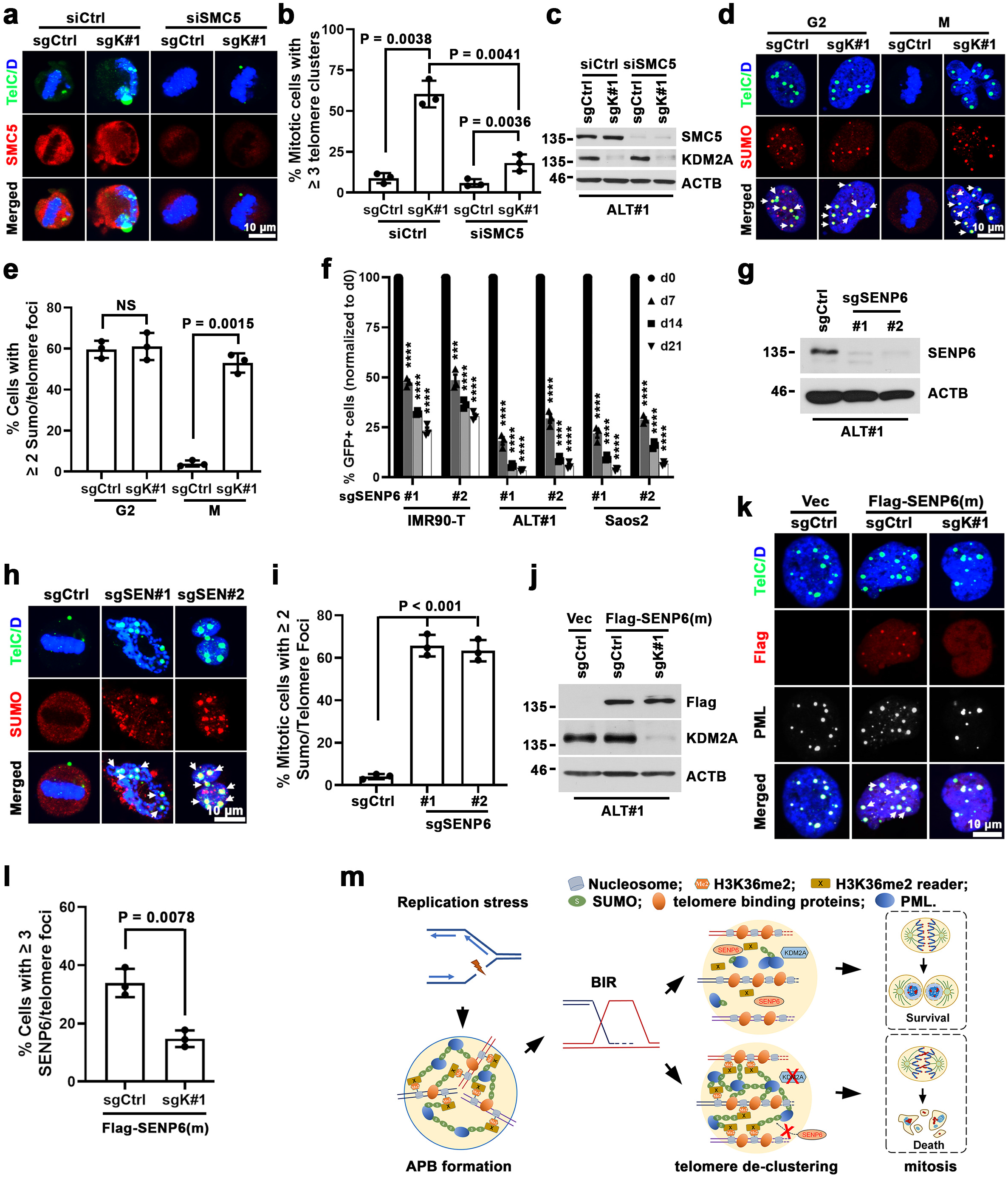
KDM2A facilitates SENP6-mediated ALT telomere de-SUMOylation. **a** Representative immuno-FISH images of multitelomere clusters in control or SMC5 siRNA transfected mitotic ALT#1 cells expressing sgCtrl or sgK#1. The mitotic (M) cells were prepared from G2 synchronized cells after 1 h release from CDK1 inhibitor. The arrows denote abnormal mitotic telomere cluster foci. **b** Percentages of mitotic cells containing ≥ 3 telomere cluster foci are expressed as means ± s.e.m. of three independent experiments, unpaired t-test. **c** Western blot analysis of SMC5 expression in whole-cell lysates prepared from the siCtrl or siSMC5-transfected ALT#1 cells. **d** Representative immuno-FISH images of telomere SUMOylation in sgCtrl or sgK#1-transduced ALT#1 cells. The mitotic (M) cells were from G2 synchronized cells upon 1 h release from CDK1 inhibitor. The arrows denote SUMO2/3-associated telomere foci. **e** Percentages of cells containing ≥ 2 SUMO-associated telomere foci. Data are expressed as means ± s.e.m. of three independent experiments, unpaired t-test. **f** Competition-based proliferation assay of the indicated *SENP6*-targeted sgRNAs (sgSEN#1 and sgSEN#2) on the fitness of IMR90-T, ALT#1 and Saos2 cells. The data are expressed as means ± s.e.m. of three independent experiments. **g** Western blot analysis of SENP6 in whole-cell lysates prepared from sgCtrl, sgSEN#1 or sgSEN#2-transduced ALT#1 cells. **h** Immuno-FISH analysis of telomere SUMOylation in mitotic ALT#1 cells transduced with sgCtrl, sgSEN#1 or sgSEN#2. The mitotic cells were from G2 synchronized cells upon 1 h release from CDK1 inhibitor. The arrows denote SUMO2/3-associated telomere foci. **i** Percentages of mitotic cells containing ≥ 2 SUMO-associated telomere foci are expressed as means ± s.e.m. of three independent experiments, unpaired t-test. **j** Western blot analysis of Flag and ACTB (loading control) in whole-cell lysates prepared from the indicated ALT#1 cells transduced with Flag-tagged SENP6^C1030A^ mutant. **k** Immuno-FISH analysis of telomere and SENP6^C1030A^ association in Flag-SENP6^C1030A^ - expressing ALT#1 cells transduced with sgCtrl or sgK#1. The arrows denote SENP6^C1030A^- associated telomere foci. **l** Percentages of cells containing ≥ 3 Flag-SENP6^C1030A^-associated telomere foci are expressed as means ± s.e.m. of three independent experiments, unpaired t-test. **m** A model illustrating the key events in ALT pathway.

ALT telomere clustering and phase-separated APB assembly are known to rely on SUMOylation of telomere associated proteins ^41, 44, 45^. Immuno-FISH analysis of G2-synchronized control and KDM2A-depleted ALT#1 cells revealed comparable levels of SUMO2/3-associated telomere foci (SUMO-T) formation (Fig. 7d, e), suggesting that KDM2A is not crucial for telomere protein SUMOylation. Upon release from the CDK1 inhibitor block, the control ALT#1 cells that had entered mitosis largely cleared their SUMO-T foci and multitelomere clusters. By contrast, we found that a large portion of sgK#1-transduced mitotic ALT#1 cells maintained their SUMOylation at the aberrantly retained multitelomere clusters (Fig. 7d, e), suggesting a role of KDM2A in regulating telomere protein de-SUMOylation. Similar results were also obtained in Saos2 cells depleted of KDM2A expression (Supplementary Fig. 14c, d), although the intensities of telomere SUMOylation in Saos2 cells were notably lower than those in ALT#1 cells.

SUMO de-conjugation from substrates is catalyzed by a group of SENP family of isopeptidases ^46^. To interrogate the role of de-SUMOylation in ALT telomere maintenance and cell survival, we assembled a focused sgRNA sub-library targeting the 7 human SENP family members (10 sgRNAs/gene) (Supplementary Table 3). The dropout screens of the isogenic ALT (ALT#1, #2 and #3) and paired IMR90-T control cells (IMR90-T#1 and #2) scored SENP6 as the only member that was differentially required by ALT cells (Supplementary Fig. 14e). GFP-based competition assays of two independent *SENP6-*sgRNAs (sgSEN#1 and #2) further confirmed its essentiality to ALT-dependent ALT#1 and Saos2 cells (Fig. 7f). Notably, depletion of SENP6 also affected the growth of IMR90-T#1 cells, although to a lesser extent than ALT#1 and Saos2 cells. To examine whether SENP6 loss interferes with the de-SUMOylation at ALT telomeres, the sgCtrl, sgSEN#1 or sgSEN#2-transduced ALT#1 cells were firstly synchronized to the G2 phase by sequential thymidine and CDK1 inhibitor treatment. Further immuno-FISH analysis of the G2-synchronized control and SENP6-depleted ALT#1 cells revealed comparable levels of SUMO-T foci formation (Supplementary Fig. 14f, g), suggesting that SENP6 is not crucial for ALT telomere protein SUMOylation or APB formation. After released from CDK1 inhibitor treatment, the control ALT#1 cells that had entered mitosis largely cleared their telomere SUMOylation and SUMO-T foci. By contrast, a large percentage of SENP6-depleted mitotic ALT#1 cells abnormally retained SUMOylated multitelomere clusters (Fig. 7g-i), reminiscent of KDM2A depletion. Similar phenotypes were also observed in Saos2 cells depleted of SENP6 expression (Supplementary Fig. 14h-l), indicating that SENP6-mediated de-SUMOylation is required for telomere cluster dissolution following ALT-directed telomere synthesis.

Since inactivation of KDM2A and SENP6 in ALT cells induced a similar telomere declustering defect, we next examined whether KDM2A loss might interfere with ALT telomere protein de-SUMOylation through blocking SENP6 recruitment to its substrates. Notably, while SENP6 generally binds to its substrates transiently, mutation of the catalytic cysteine to alanine (C1030A) stabilizes the interaction ^47^. To test whether SENP6 physically interacts with ALT telomeres, we expressed the *SENP6*^*C1030A*^ mutant transgene in ALT#1 cells (Fig. 7j). Analysis of the G2-synchronized ALT#1 cells found that the Flag-tagged SENP6^C1030A^ mutant was indeed localized to the clustered ALT telomeres (Fig. 7k). Importantly, quantitation of the telomere-associated SENP6^C1030A^ foci revealed that KDM2A depletion significantly diminished the recruitment of SENP6^C1030A^ to telomeres (Fig. 7l). Similar results were also obtained in Saos2 cells depleted of KDM2A expression (Supplementary Fig. 14m-o), suggesting that KDM2A may function to promote ALT telomere de-clustering by facilitating SENP6-mediated telomere de-SUMOylation. Consistently, complementation of CRISPR-resistant cDNAs encoding wild-type, but not KDM2A mutant defective of demethylase (D214A) or chromatin binding (S603D), restored SENP6^C1030A^ recruitment to ALT telomeres in sgK#1-transduced ALT#1 cells (Supplementary Fig. 15a-c). Finally, it is worthy to note that our co-immunoprecipitation analysis uncovered no evidence of physical interaction between KDM2A and SENP6 (Supplementary Fig. 16), suggesting that KDM2A may regulate SENP6 recruitment to ALT telomeres indirectly through modulating telomere H3K36 methylation.

## Discussion

The implementation of the genetic concept of synthetic lethal or synthetic lethal-like interaction holds great promise in anticancer target discovery. By undertaking chromatin regulator-focused genetic screens in a well-controlled isogenic ALT cellular model system, we identified histone demethylase KDM2A as a selective molecular vulnerability of ALT-dependent cancers. We further demonstrate that KDM2A is required for post-recombination ALT telomere declustering. We show that depletion of KDM2A in ALT cells impairs isopeptidase SENP6-mediated SUMO deconjugation at ALT telomeres. Inactivation of KDM2A or SENP6 compromises post-recombination ALT multitelomere de-SUMOylation and dissolution that subsequently lead to chromosome missegregation and mitotic cell death. Results from this study thus support efforts to develop KDM2A inhibitors targeting ALT cancers.

Synthetic lethality provides a framework for targeting loss of function of tumor suppressor and DNA repair genes, as exemplified by the success of PARP inhibitors in the treatment of BRCA1/2-deficient tumors ^48, 49^. The advance of CRISPR-based screen methodology has further enabled large-scale studies to profile synthetic lethal or synthetic lethal-like interactions in a diverse collection of human cancer cells across many genetic contexts ^50-52^. But despite the fact that ALT is utilized in a substantial fraction (5% - 10%) of human cancers, ALT cancer cells were infrequently included in those screening initiatives. The poor representation is likely due to the paucity of ALT cancer cell lines suitable for large-scale functional genomic screens. To overcome this hurdle, we employed an *in vitro* ALT-immortalized isogenic model system. The inclusion of isogenic non-ALT derivatives in our screens greatly reduced potential confounding variables such as cell type difference and co-occurring genetic changes, making it easy to infer the genetic interactions from a small panel of paired cell lines. Our study thus demonstrates the applicability of well-controlled isogenic models in identifying and prioritizing targets from genetic interaction screens.

Ideal anticancer therapies should demonstrate a robust therapeutic window toward tumor cells by sparing normal cells and limiting off-tumor toxicity. This study identifies KDM2A as a promising therapeutic target that is selective for ALT-dependent cancers. Inactivation of KDM2A in ALT cells causes robust cell growth inhibition by inducing chromosome missegregation and mitotic cell death. By comparison, KDM2A is largely dispensable for non-ALT cell growth. Our competition-based proliferation and total knock-out assays of a panel of non-ALT cells of diverse tissue origins found that KDM2A inactivation leads to a minor reduction in cell proliferation without significantly affecting cell survival. Consistently, conditional *Kdm2a* deletions in the mouse myeloid compartment revealed no discernible impact on the development and maturation of myeloid cells ^53^. These findings suggest a potentially large therapeutic window for future KDM2A-targeted therapies for ALT cancer treatment.

Emerging evidence suggests that ALT-directed telomere elongation emanates from telomere replication stress and proceeds through a conservative BIR-like pathway ^5, 6, 20, 22, 41^. However, the molecular signals that control ALT pathway initiation and termination still remain largely unclear. For example, although APB formation is known to depend on protein SUMOylation ^45^, how telomere replication stress signals proceed to promote APB formation in ALT cells is poorly understood. Equally unclear are the molecular events that govern the dissolution of the clustered ALT telomeres after completion of the ALT-directed telomere DNA synthesis. In this study, we demonstrate that post-recombination ALT telomere de-clustering requires SENP6-mediated de-SUMOylation. Our data indicate that this process is regulated by KDM2A-directed telomere histone H3K36me2 demethylation, underscoring its importance in ALT-directed telomere maintenance.

Histone methylation pathways play important roles in orchestrating DNA damage signaling ^54-56^. Among the total of 21 histone lysine demethylases, our genetic screen identifies KDM2A as the only one that is selectively essential for ALT cell growth. Consistent with a previous proteomic study ^33^, our telomere dot-blot assay of chromatin immunoprecipitation demonstrates the physical interaction of KDM2A with ALT telomeres, supporting a direct KDM2A involvement in ALT-dependent telomere maintenance. KDM2A is a lysine-specific demethylase that targets lower methylation states of H3K36 (Kme1 and Kme2) ^25, 26^. Indeed, depletion of KDM2A in ALT cells leads to increased H3K36me2 at telomeres. But interestingly, KDM2A is not required for ALT-directed telomere DNA synthesis and APB formation. Instead, we found that KDM2A-mediated demethylation promotes post-recombination ALT telomere declustering, at least partly through facilitating SENP6-mediated telomere de-SUMOylation. Given that H3K36 methylation is a major event following DNA double strand break (DSB) induction to recruit DNA repair proteins ^57, 58^, we propose a model in which KDM2A functions as an epigenetic eraser of H3K36me2-dependent signaling that safeguards proper chromosome segregation by facilitating SENP6-mediated ALT telomere de-SUMOylation and declustering (Fig. 7m). Loss of KDM2A impairs SENP6 recruitment to ALT telomeres and thus their declustering, leading to chromosome missegregation and mitotic death. But despite our data that support a direct KDM2A involvement in ALT-mediated telomere maintenance, we cannot exclude the possibility that KDM2A may contribute indirectly through transcriptional regulation of genes involved in the process. Also noticeably, our co-immunoprecipitation analysis did not find evidence of physical KDM2A and SENP6 interaction. Future studies are needed to sort out the detailed molecular events that follow KDM2A action.

Our study nominates KDM2A as a selective molecular vulnerability of ALT-dependent cancer cells. In human cancers, ALT activation is strongly associated with mutations of the chromatin modulator genes *ATRX* and *DAXX* ^9-13^. But notably, our current study reveals that KDM2A is not a simple synthetic lethal vulnerability with *ATRX* or *DAXX* loss. Instead, we found that KDM2A is critical for ALT-directed telomere maintenance, which is utilized by cells deficient of ATRX or DAXX. These findings suggest that KDM2A-mediated demethylation may play an important role in resolving the telomere replication stress and repair-associated intermediate structures following homology-directed telomere synthesis. As replication stress-induced DNA damage and repair are common features of human cancers, it will be interesting in the future to evaluate the molecular functions of KDM2A and, more broadly, H3K36 methylation and demethylation in those processes.

## Methods

### Cell lines and plasmids

The human cell lines A172, G292, HEK293T, HeLa, Hs792, IMR90, MCF-7, MG63, NCI-H1299, U118, and U2OS were obtained from ATCC. The IMR90-T, ALT#1, #2 and #3 cells were derived from large T-transformed IMR90 cells as previously described ^22^. The patient-derived *ATRX*-mutant glioblastoma cell line pGBM6 was established from collected tumor specimens after obtaining written informed consent preoperatively and approved by the Institutional Reviewer Boards of the Southwest Hospital. The human glioblastoma cell line LN464 was kindly provided by F. Furnari (University of California, San Diego). The clonally derived *KDM2A*-knockout HeLa and LN464 cell lines, or *ATRX*-depleted IMR90-T and LN464-T cells were generated using lentiviruses produced in HEK293T cells with lentiCRISPR-v2 vectors containing *KDM2A* or *ATRX* targeting sgRNA and selected with blasticidin as previously described ^22^. For the cDNA expression experiments, full length cDNAs was cloned into a lentiviral expression vector pLU-IRES-Puro, -Blast or -Neo vector containing 3xN terminal Flag or C-terminal GFP tag. CRISPR sgRNA resistant synonymous or functional domain point mutations were introduced by PCR mutagenesis using NEBuilder HiFi DNA Assembly Master Mix (NEB, E2631). Stable cell lines were generated using the pLU vectors and selected with puromycin or blasticidin. Cell lines obtained were not authenticated and were tested negative for mycoplasma. In this study, all cells were cultured with respective media in a humidified 37°C, 5% CO2 incubator. All the sgRNAs targeting human genes were cloned into lentiCRISPR v2 (Addgene, #52961), lentiCas9-Blast (Addgene, #52962), LRG2.1 (U6-sgRNA-GFP, Addgene, #108098) or LRPuro (U6-sgRNA-Puromycin) as indicated. Single sgRNAs were cloned by annealing two DNA oligos and ligating into a BsmB1-digested vector. To improve U6 promoter transcription efficiency, an additional 5’ G nucleotide was added to all sgRNA oligo designs that did not already start with a 5’ G. A list of sgRNA information is provided in Supplementary Table 4.

### Animal experiments

NSG (NOD.Cg-PrkdcscidIl2rgtm1Wjl/SzJ) mice were purchased from Jackson laboratories. All of mouse use and procedures were approved by the Institutional Animal Care and Use Committee of the Weill Cornell Medicine. Mice were maintained on a 12 h light/dark cycle, and food and water was provided ad libitum. For all mouse studies, mice of either sex were used, and mice were randomly allocated to experimental groups, but blinding was not performed. Mice (aged 5–8 weeks) were age-matched for tumor inoculation. Group sizes were selected on the basis of prior knowledge. For subcutaneous grafting, sgCtrl, sgK#1 or #2 transduced Saos2 cells were resuspended in 50% Matrigel (BD Bioscience, #356231) in PBS and ~5,000,000 cells were injected into each flank of NSG mice. Tumor growth was monitored and measured every 7 days by caliper, and volume was calculated by the formula: 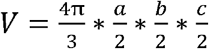 (V tumor volume; a tumor length; b tumor width; c tumor height). The end points were determined on the basis of the level of animal discomfort and tumor sizes.

### Construction of pooled sgRNA library

A gene list of 455 chromatin modification-associated factors in the human genome was manually curated. 4-14 sgRNA were designed against functional domains of each gene based on the domain sequence information retrieved from NCBI Conserved Domains Database. The design principle of sgRNA was based on previous reports and the sgRNAs with the predicted high off-target effect were excluded ^59-61^. All of the sgRNAs oligos including positive and negative control sgRNAs were synthesized in a pooled format (CustomArray Inc) and PCR amplified. The library was constructed by cloning the PCR amplified products into BsmB1-digested LRPuro vector. The identity and relative representation of sgRNAs in the pooled plasmids were verified by a deep-sequencing-based analysis.

### Construction of CRISPR-based KDM2A exon-tiling library

A list of total 492 sgRNAs that covered the entire open reading frame of *KDM2A* was designed by excluding the ones with predicted high off-target effect ^59-61^. All of the sgRNAs oligos including positive and negative control sgRNAs were synthesized in a pooled format (CustomArray Inc). The library was generated by cloning the PCR amplified products of the synthesized sgRNA oligos into BsmB1-digested LRPuro vector. The identity and relative representation of sgRNAs in the pooled plasmids were verified by a deep-sequencing-based analysis.

### Lentiviral transduction

Lentiviruses were produced by co-transfection of indicated plasmids and packaging vectors into HEK293T packaging cells as previously described ^22^. In brief, to generate lentivirus, 8 × 10^6^ 293T cells in 100□mm tissue culture dishes were transfected with mixture of 8.5□μg of plasmid DNA, 4 μg of pMD2.G and 6 μg of psPAX2 packaging vectors, and 45 μl of 1 mg/mL Polyethylenimine (PEI 25000). The media was replaced 6-8 h post transfection. The virus-containing supernatant were collected at 48 and 72□h post transfection and pooled. For infection, virus-containing supernatant was mixed with the indicated cell lines supplied with 4 mg/mL polybrene, and then centrifuged at 2,000 rpm for 30 min at room temperature. Fresh media was changed 24 h post-infection. Antibiotics (10 μg/mL blasticidin, 2 μg/mL puromycin, 500 μg/mL G418, and/or 200 μg/mL hygromycin) were added 48 h post-infection when selection was required.

### Pooled CRISPR-Cas9 and KDM2A exon-tilling screen

The pooled CRISPR-based negative selection and KDM2A exon-tiling screens were carried out as previously described with some modifications ^62^. In brief, Cas9-expressing cells were infected with the lentiviral chromatin modification-associated factor- or KDM2A exon-tiling sgRNA library at an MOI 0.3 – 0.4 such that every sgRNA is represented in ~ 1,000 cells. Fresh media was changed 24 h post-infection. At 48 h post infection, puromycin (2 μg/mL) was added and the infected cells were selected with for 48–72 h. To maintain the representation of sgRNAs, the number of infected cells was kept at least 1000 times the sgRNA number in the library during the screen. Start-point cells were harvested at day 5 post-infection and served as a reference representation of the pooled sgRNA library. Cells were cultured for 16 population doublings and harvested as the end time point. Genomic DNA was extracted from cell pellets using QIAamp DNA Blood Maxi Kit (QIAGEN, #51194). The sgRNA cassette was PCR amplified from genomic DNA using Phusion High-Fidelity PCR Master Mix (New England Biolabs, M0531S). The amplified products were pooled and amplified again via PCR using primers harboring Illumina TruSeq adapters with i5 and i7 barcodes, and the resulting libraries were sequenced on an Illumina Nextseq500. The sequencing data was de-multiplexed. The read count of each sgRNA was calculated with no mismatches by comparing to the reference sequence. Individual sgRNAs with the read count lower than 50 in the initial time point were discarded.

### Competition-based proliferation assay

The flow cytometry analysis (FACS)-based dropout assay was performed as previously described ^22^. In brief, the indicated Cas9-expressing cell lines were transduced with a lentiviral LRG2.1 (U6-sgRNA-GFP, Addgene, #108098) sgRNA vector that co-expresses a GFP reporter. The percentage of sgRNA transduced GFP-positive cell population in culture was measured at indicated time points using a LSRII flow cytometer (BD Biosciences). The change of GFP percentage was used to assess proliferation of sgRNA-transduced cells relative to the non-transduced cells in the culture.

### Clonogenic-based proliferation assays

To evaluate growth rate difference, indicated control and experimental cells were plated at 10,000 - 20,000 cells per well of 6-well plate. Cells were cultured with respective media in a humidified 37°C, 5% CO2 incubator for 15 – 25 days before fixed with 10% formalin and stained with 0.1% crystal violet.

### Immuno-FISH assay

Indirect immunofluorescence (IF) combined with fluorescence *in situ* hybridization (FISH) analysis was performed as previously described ^22^. Briefly, cells grown on coverslips were fixed for 15 min in 4% paraformaldehyde at RT, followed by permeabilization for 10 min in PBS with 0.3%Triton X-100 at RT. After washing with 1x PBS, cells were incubated for 60 min in blocking solution (1% BSA, 10% FBS, 0.2% fish gelatin, 0.1% Triton X-100, 1 mM EDTA in 1x PBS) before immuno-staining. Primary antibodies were prepared in blocking solution as following dilutions: 53BP1 (1:500, IHC-00001; Bethyl Laboratories), 53BP1 (1:300, AF1877; R&D Systems), BLM (1:250, Cat# A300-110A; Bethyl Laboratories), FLAG (1:200, F1804; Sigma), γH2AX (1:2,000, A300-081A; Bethyl Laboratories), phospho-Histone H3 (Ser10) (1:200, #9701; Cell Signaling), PML (1:200, sc-966; Santa Cruz Biotechnology), PML (1:500, ab96051; abcam), SMC5 (1:400, A300-236A; Bethyl Laboratories), SUMO-2/3 (1:250, ab3742; abcam), and TRF1 (1:200, ab10579; abcam). After incubated with indicated primary antibodies at RT for 2 h, cells were washed 3 times with PBST (1xPBS containing 0.1% Tween-20) before incubated with indicated secondary antibodies conjugated to fluorophores diluted in the same solution for 45 min at RT. After 3 washes with PBST, cells were fixed again with PFA for 10 min, then washed in 1x PBS, dehydrated in ethanol series (70%, 95%, 100%), and air-dried. Coverslips were denatured for 10 min at 85°C in hybridization mix [70% formamide, 10 mM Tris-HCl, pH 7.2, and 0.5% blocking solution (Roche)] containing 100nM telomeric PNA probe TelC-FITC (F1009, PNA Bio), and hybridization was continued at RT for 2 h in the dark moisturized chambers. Coverslips were washed three times with Wash solution (70% formamide, 2x SSC) for 10 min each and in 2x SSC, 0.1% tween-20 three times for 5 min each. During the second wash, cells were stained with DAPI. Slides were mounted with VectorShield (Vector Laboratories). Images were captured with a 60x lens on an Olympus FLUOVIEW laser scanning confocal microscope (Olympus).

### Protein extraction and western blotting

Cells were lysed in RIPA buffer (150 mM NaCl, 50 mM Tris, 0.5% Na-Deoxycholate, 0.1% SDS and 1% NP-40), and equal amount of protein was resolved using Nupage Novex 4-12% Bis-Tris Gel (ThermoFisher Scientific) as previously described ^22^. Primary antibodies used were β-ACTIN (ACTB) (1:5,000, #2228; Sigma), ATRX (1:1,000, sc-15408; Santa Cruz Biotechnology), FLAG (1:1,000, F1804; Sigma), H3 (1:2,000, ab1791; abcam), H3K36me2 (1:1,000, 2901; Cell Signaling), H3K36me3 (1:2,000, 61101; Active Motif), KDM2A (1:1,000, A301-476A; Bethyl Laboratories), SENP6 (1:500, HPA024376; Sigma-Aldrich), SMC5 (1:2000, A300-236A; Bethyl Laboratories), and TUBULIN (1:2,000, ab15246; abcam). Secondary antibodies used were donkey anti-rabbit HRP (1:1,000, sc-2077; Santa Cruz Biotechnology), donkey anti-mouse HRP (1:1,000; sc-2096, Santa Cruz Biotechnology) and donkey anti-goat HRP (1:1,000, sc-2056, Santa Cruz Biotechnology). Antibody signal was detected using the ECL Western Blotting Substrate (W1015, Promega) and X-ray film (F-BX810, Phenix).

#### Cell cycle synchronization

Cells were synchronized in G2 phase by sequential treatment of thymidine and CDK1 inhibitor Ro-3306 (S7747, Selleckchem). Briefly, cells were first cultured in medium containing 2mM thymidine (Sigma-Aldrich) for 21 h. After washing twice with PBS followed by once with growth media, the cells were released into fresh medium for 3 h before treatment with 10mM CDK1 inhibitor for 12 h.

### Flow cytometry analysis (FACS)

Cells collected by trypsin and re-suspended in PBS containing 1 mM EDTA were fixed in ice-cold ethanol overnight. Ethanol-fixed single-cell suspensions were stained for DNA analysis with 1% BSA, 0.1% Tween-20, 0.1 mM EDTA, 0.5 mg/mL RNaseA, 10 µg/mL propidium iodide (PI) in 0.25 ml PBS. Cells were incubated for 30 min at 37 °C and equilibrated at room temperature in the dark for at least 10 min. Cells were analyzed by a LSRII flow cytometer (BD Biosciences). Cell cycle population analysis was conducted with FlowJo v5 software (FlowJo).

### Terminal restriction fragment (TRF) analysis

TRF analysis was conducted as previously described 22. In brief, genomic DNA was prepared using Wizard genomic DNA purification kit (Promega) as manufacturer’s instruction. For telomere length and Southern blot analysis, genomic DNA (~ 5 μg) was digested with *Alu*I+*Mbo*I restriction endonucleases, fractionated in a 0.7% agarose gel, denatured, and transferred onto a GeneScreen Plus hybridization membrane (PerkinElmer). The membrane was cross-linked, hybridized at 42°C with 5′-end-labeled ^32^P- (TTAGGG)_4_ probe in Church buffer, and washed twice for 5 min each with 0.2 M wash buffer (0.2 M Na_2_HPO4 pH 7.2, 1 mM EDTA, and 2% SDS) at room temperature and once for 10 min with 0.1 M wash buffer at 42°C. The images were analyzed by Phosphor-imager, visualized by Typhoon 9410 Imager (GE Healthcare), and processed with ImageQuant 5.2 software (Molecular Dynamics).

### C-circle assay

C-circle assay was performed as described ^21, 63^ with minor modifications. Briefly, genomic DNA digested with *Alu*I and *Mbo*I was cleaned up by phenol-chloroform extraction and precipitation. An aliquot of purified DNA was diluted in nuclease-free water, and concentrations were measured to the indicated quantity (~15 ng/µl) using a Nanodrop spectrophotometer (Thermal Scientific). Sample DNA (30 ng in a total volume of 10 μl) was combined with 10 μl reaction mix [0.2 mg/ml BSA, 0.1% Tween, 0.2 mM each dATP, dGTP, dTTP, 2 × □29 Buffer, and 7.5 U □DNA polymerase (NEB)]. The reactions were mixed well, incubated at 30°C for 8 h, and then at 65°C for 20 min. The reaction products were diluted to 400 μl with 2 × SSC, dot-blotted onto a 2 × SSC-soaked GeneScreen Plus membrane, and hybridized with a ^32^P-labeled (CCCTAA)_4_ probe at 37°C for overnight to detect C-circle amplification products. The blots were washed four times at 37°C in 0.5 × SSC/0.1% SDS buffer, exposed to Phosphor-imager screens, visualized by Typhoon 9410 Imager (GE Healthcare Life Sciences), and quantified with ImageQuant 5.2 software (Molecular Dynamics).

### Chromatin immunoprecipitation (ChIP) assays

ChIP assays were performed as described previously ^22^. Briefly, cells (~ 1 × 10^7^) were crosslinked in 1% formaldehyde with shaking for 15 mins, quenched by the addition of glycine to a final concentration of 0.125 M, and lysed in 1 ml SDS lysis buffer (1% SDS, 10 mM EDTA, and 50 mM Tris-HCl, pH 8.0) supplemented with 1 mM PMSF and protease inhibitor cocktails (Sigma-Aldrich). The lysates were sonicated with a Diagenode Bioruptor, cleared by centrifugation to remove insoluble materials, and diluted 10 fold into IP Buffer (0.01% SDS, 1.1% Triton X-100, 1.2 mM EDTA, 16.7 mM Tris pH 8.1, 167 mM NaCl, 1 mM PNSF, and protease inhibitors cocktail) for IP reaction at 4°C overnight. Each immuno-complex was washed five times (1 ml wash, 10 mins each) in ChIP related wash buffer at 4°C, eluted by addition of 150µl Elution buffer (10 mM Tris, pH 8.0, 5 mM EDTA, and 1% SDS) at 65°C for 30 mins, and the elutes were placed at 65°C for overnight to reverse cross-linking. The elutes was further treated with Proteinase K in a final concentration of 100 µg/ml at 50°C for 2 h, and ChIP DNA was purified by Quick PCR Purification Kit (Life Technologies) following the manufacturer’s instruction. ChIP DNA was denatured, dot blotted onto GeneScreen Plus blotting membranes (PerkinElmer) and crosslinked at 125 mJ. Oligonucleotide probe for telomere or Alu repeats was labeled with [^32^P]-ATP (3,000 Ci/mmol) and T4 nucleotide kinase (New England Biolabs). The membrane was hybridized in Church hybridization buffer containing a ^32^P-labeled probe at 42°C overnight, washed three times in 0.04 N Na-phosphate, 1% SDS, 1 mM EDTA at 42°C, developed with a Typhoon 9410 Imager (GE Healthcare Life Sciences) and quantified with ImageQuant 5.2 software (Molecular Dynamics). Antibodies used in ChIP assay were anti-FLAG (F1804, Sigma), anti-H3K36me2 (2901, Cell Signaling), and mouse IgG (sc2025, Santa Cruz Biotechnology).

### Detection of ALT-directed telomere DNA synthesis

To visualize DNA synthesis at telomeres, synchronized G2 cells were incubated with 20 mM EdU for 2 h. Cells were permeabilized, then fixed with 4% formaldehyde PBS solution. The Click-iT® Alexa Fluor 555 azide reaction was then performed according to the manufacturer’s instructions (Click Chemistry Tools). After washing with 1x PBS, cells were incubated for 60 min in blocking solution (1% BSA, 10% FBS, 0.2% fish gelatin, 0.1% Triton X-100, 1 mM EDTA in 1x PBS) before PML immuno-staining at RT for 2 h. After incubation of secondary antibodies and 3 washes with PBST, cells were fixed again with PFA for 10 min at RT, washed in 1x PBS, dehydrated in ethanol series (70%, 95%, 100%), and air-dried. Coverslips were denatured for 10 min at 85°C in hybridization mix [70% formamide, 10 mM Tris-HCl, pH 7.2, and 0.5% blocking solution (Roche)] containing 100nM telomeric PNA probe TelC-FITC (F1009, PNA Bio), and hybridization was continued for 2 h at room temperature in the dark moisturized chambers. Coverslips were washed three times with Wash solution (70% formamide, 2x SSC) for 10 min each and in 2x SSC, 0.1% tween-20 three times for 5 min each. During the second wash, cells were stained with DAPI. Slides were mounted with VectorShield (Vector Laboratories). Images were captured with a 60x lens on an Olympus FLUOVIEW laser scanning confocal microscope (Olympus).

### Live cell imaging

H2B-GFP-expressing control or KDM2A targeting sgRNA transduced cells were seeded at 8 × 10^4^ cells per well in fibronectin coated 12-well 1.5 mm glassbottom wells (Cellvis). Following culture in DMEM medium supplemented with 15%FBS for 24 hours, cells were synchronized in G2 phase by sequential treatment of thymidine and CDK1 inhibitor Ro-3306 (S7747, Selleckchem). Cells were firstly cultured in medium containing 2mM thymidine (Sigma-Aldrich) for 24 h. After washing twice with PBS followed by once with growth media, the cells were then released into fresh medium for 2 h before treatment with 10mM CDK1 inhibitor for 16 h. Finally, cells were washed twice with PBS and once with growth media before subjected to live cell imaging with Zeiss Cell Observer (ZEISS). Cells were monitored for 8 hours at 5 min intervals. Movies are output by Zeiss ZEN software (ZEISS).

### Statistical Analysis

Details regarding quantitation and statistical analysis are provided in the figures and figure legends. We determined experimental sample sizes on the basis of preliminary data. All results are expressed as mean ± s.e.m. GraphPad Prism software (version 7.0e) was used for all statistical analysis. Normal distribution of the sample sets was determined before applying Student’s two-tailed t test for two group comparisons. Differences were considered significant when P < 0.05.

## Data availability

The data that support the findings of this study are available within the article and its Supplementary Information files. All the uncropped western blots and raw data are provided as a Source data file.

## Code availability

For specific requests, please contact the corresponding authors.

## Acknowledgements

This work was partly supported by a research award from the William Rhodes and Louise Tilzer-Rhodes Center for Glioblastoma at NewYork-Presbyterian Hospital. Additional funding was provided to J. Paik by NIH/NCI grant P01 #CA214274. J-Y. Jang is partly supported by Basic Science Research Program through the National Research Foundation of Korea (NRF) funded by the Ministry of Education (2021R1A6A3A03039136). H. Zheng was partly supported by the Sontag Foundation and the Chen & Xiao Anti-Cancer Foundation.

## Author contributions

FL, YW, IH, JYJ, LX, ZD, EYY, YC, CW, ZH, YHH, XH, and LZ carried out experiments. YJ performed the sequencing data analysis. NFL, PML, HY, JP, and HZ supervised this study. FL, JP, and HZ analyzed the data and wrote the manuscript.

## Competing interests

The authors declare no competing interests.

## Materials & Correspondence

Further information and requests for resources and reagents should be addressed to J.P. or H.Z.

## Notes

### Competing Interest Statement

The authors have declared no competing interest.

