## Supplemental Figures for "Histone demethylase KDM2A is a selective vulnerability of cancers relying on alternative telomere maintenance"

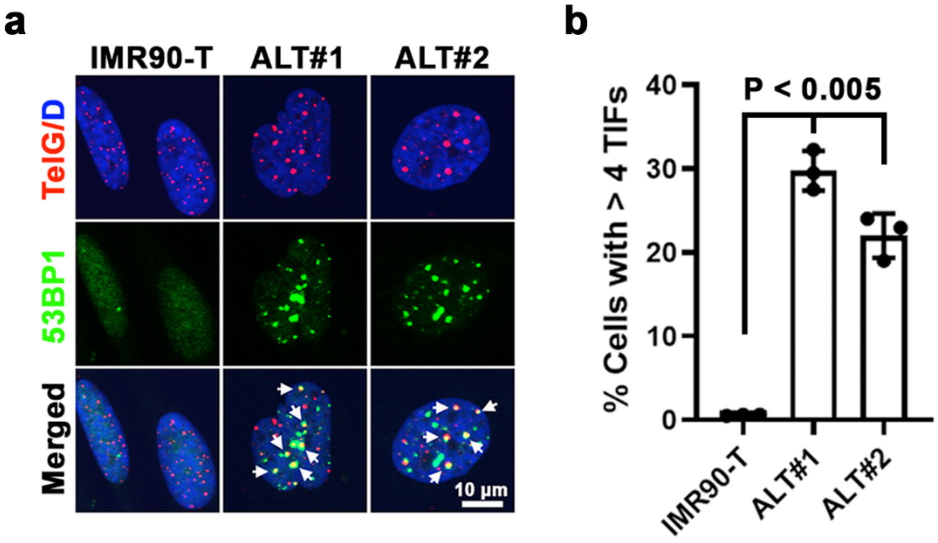


**Supplementary Figure 1. ALT-immortalized IMR90 cells exhibit high levels of telomere replication dysfunction.** **a** Representative immuno-FISH images of 53BP1 and telomeres in telomerase expressing (IMR90-T) or ALT-immortalized (ALT#1 and ALT#2) IMR90 cells. The arrows denote 53BP1-associated telomere dysfunction-induced foci (TIFs). Scale bar = 10 µm. **b** Percentages of cells containing ≥ 4 TIFs are expressed as means ± s.e.m. of three independent experiments; unpaired t-test.

Supplementary Figure 2

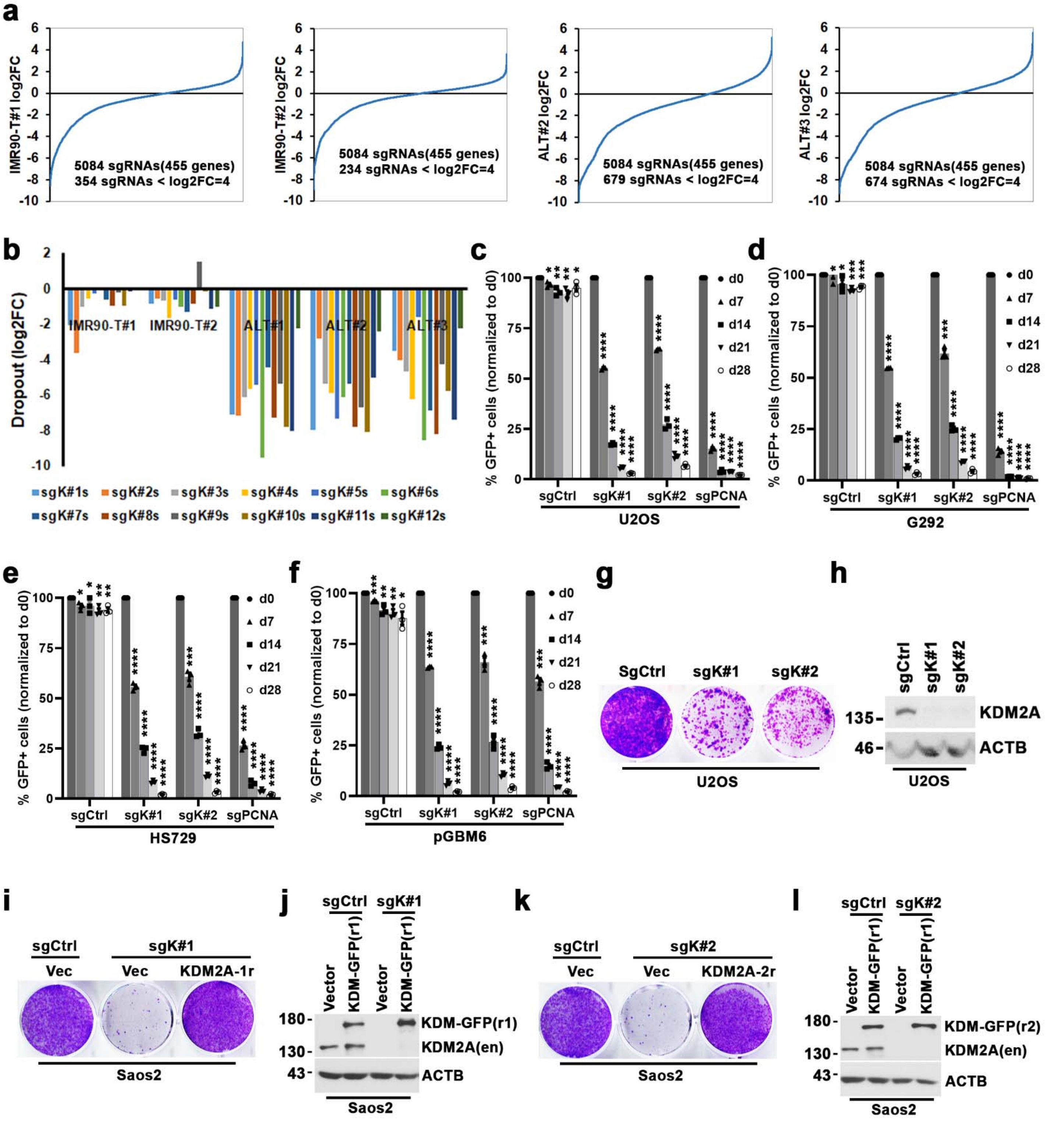


**Supplementary Figure 2. KDM2A is required for ALT-dependent cell proliferation. a** Ranking of sgRNAs by log2 fold-change (log2FC) of abundance (ratio of start to end point) in IMR90-T#1, IMR90-T#2, ALT#2, and ALT#3 cells. x axis shows targeting sgRNAs; y axis shows the log2FC of each targeting sgRNA after 16 population doublings. **b** The log2FC dropout of the abundance of individual KDM2A-targeted sgRNAs (sgK#1 - 12) in IMR90-T#1, IMR90-T#2, ALT#1, ALT#2, and ALT#3 cells after 16 population doublings. **c-f** Competition-based proliferation assay of KDM2A-targeted sgRNAs (sgK#1 and sgK#2) on the fitness of ALT+ U2OS (**c**), G292 (**d**), Hs729 (**e**), and patient-derived pGBM6 glioblastoma cells (**f**). A GFP reporter is linked to sgRNA expression. Plotted is the %GFP cells at the indicated time-points after normalized to the starting day d0 measurement. Data are expressed as means ± s.e.m. of three independent experiments; paired t-test. * denotes ns (not significant); ** $P < 0.05$; *** $P < 0.01$; **** $P < 0.001$. **g** Clonogenic assay of KDM2A-targeted sgK#1 and sgK#2 in U2OS cells. The non-targeting sgCtrl was included as a negative control. Crystal violet staining was conducted at day 15 post seeding. **h** Western blot analysis of KDM2A and ACTB (loading control) in whole-cell lysates prepared from sgCtrl, sgK#1 or sgK#2-transduced U2OS cells. **i** Clonogenic assay of sgK#1-transduced Saos2 cells expressing with control or CRISPR-resistant GFP-tagged *KDM2A* cDNA (KDM-GFP-r1). Crystal violet staining was conducted at day 22 post seeding. **j** Western blot analysis of KDM2A and ACTB in whole-cell lysates prepared from sgCtrl or sgK#1-transduced Saos2 cells expressing vector control or KDM-GFP-r1. En, endogenous KDM2A. **k** Clonogenic survival assay of sgK#2-transdued Saos2 cells expressing vector control or CRISPR-resistant *KDM2A* cDNA (KDM-GFP-r2). **l** Western blot analysis of KDM2A in the whole-cell lysates prepared from the indicated Saos2 cells.

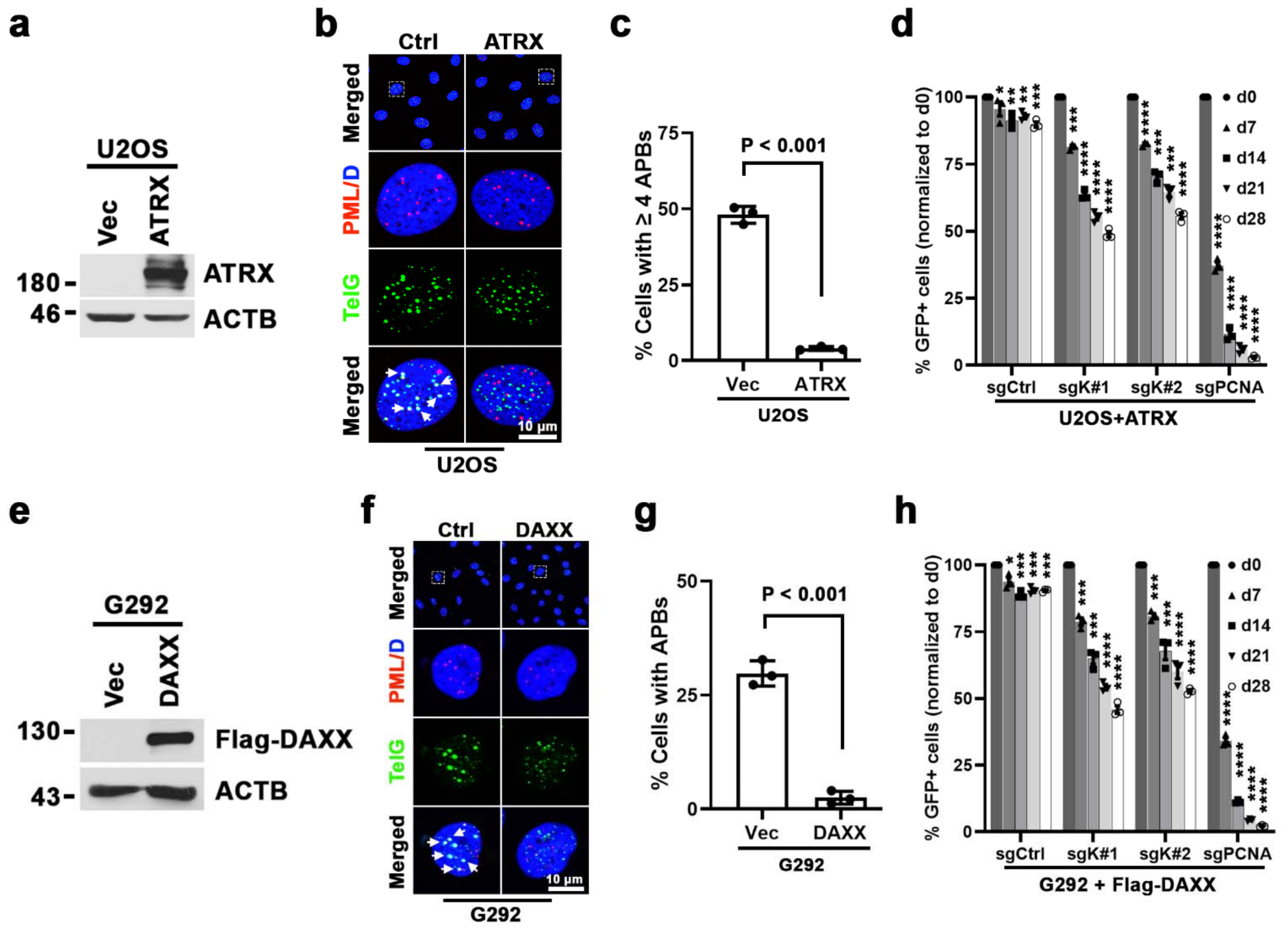


**Supplementary Figure 3. Suppression of ALT activity in ALT+ cells attenuates their KDM2A dependency. a** Western blot analysis of ATRX and ACTB in whole-cell lysates prepared from vector control or *ATRX* cDNA-transduced U2OS cells. **b** Representative immuno-FISH images of ALT-associated PML bodies (APBs) in control or ATRX-transduced U2OS cells. Scale bar = 10 µm. **c** Percentages of cells containing ≥ 4 APBs are expressed as means ± s.e.m. of three independent experiments; unpaired t-test. **d** Competition-based proliferation assay of ATRX-transduced U2OS cells. sgCtrl and sgPCNA were included as a negative or a positive control respectively. **e** Western blot analysis of Flag and ACTB in whole-cell lysates prepared from vector control or Flag-DAXX-transduced G292 cells. **f** Representative immuno-FISH images of APB formation in control or Flag-DAXX-transduced G292 cells. Scale bar = 10 µm. **g** Percentages of cells containing ≥ 4 APBs are expressed as means ± s.e.m. of three independent experiments; unpaired t-test. **h** Competition-based proliferation assay of Flag-DAXX-transduced G292 cells. In **d**, **h**, data are expressed as means ± s.e.m. of three independent experiments; paired t-test. * denotes ns (not significant); ** P < 0.05; *** P < 0.01; **** P < 0.001.

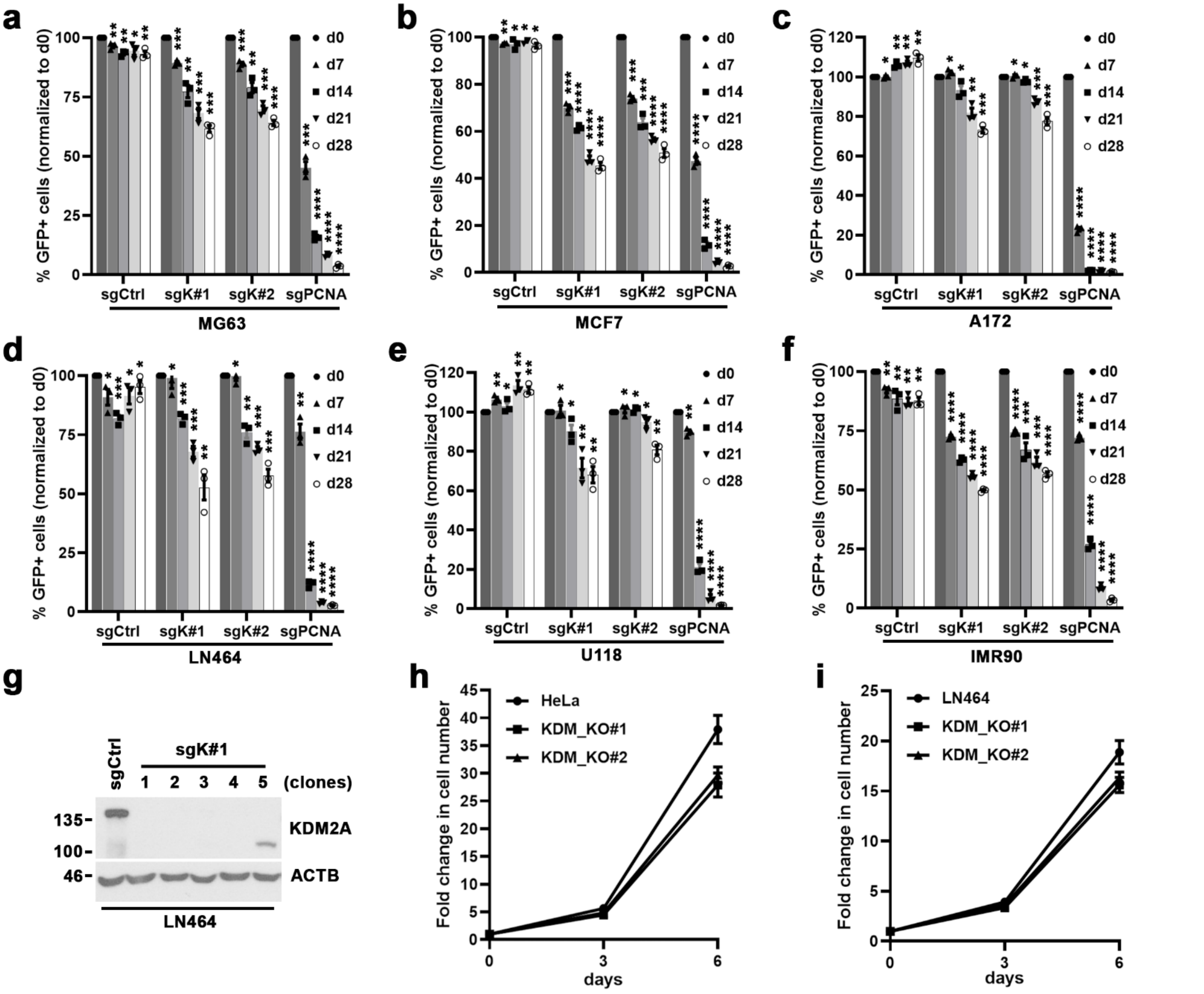

**Supplementary Figure 4. KDM2A is not essential for non-ALT cell proliferation and survival. a-f** Competition-based proliferation assay of the indicated sgRNAs in MG63 (**a**), MCF7 (**b**), A172 (**c**), LN464 (**d**), U118 (**e**), and IMR90 cells (**f**). sgCtrl and sgPCNA were included as a negative or positive control respectively. Data are expressed as means ± s.e.m. of three independent experiments; paired t-test. * denotes ns (not significant); ** $P < 0.05$; *** $P < 0.01$; **** $P < 0.001$. **g** Western blot analysis of KDM2A and ACTB (loading control) in whole-cell lysates prepared from the indicated cells. KDM2A depleted lines (clone#1-7) were established from clonally isolated LN464 cells transduced with sgK#1. The non-targeting sgCtrl transduced LN464 line was included as control. **h** Cell proliferation of wild-type or KDM2A knockout (KDM_KO#1 and #2) HeLa cells. **i** Cell proliferation of wild-type or KDM2A knockout (KDM_KO#1 and #2) LN464 cells.

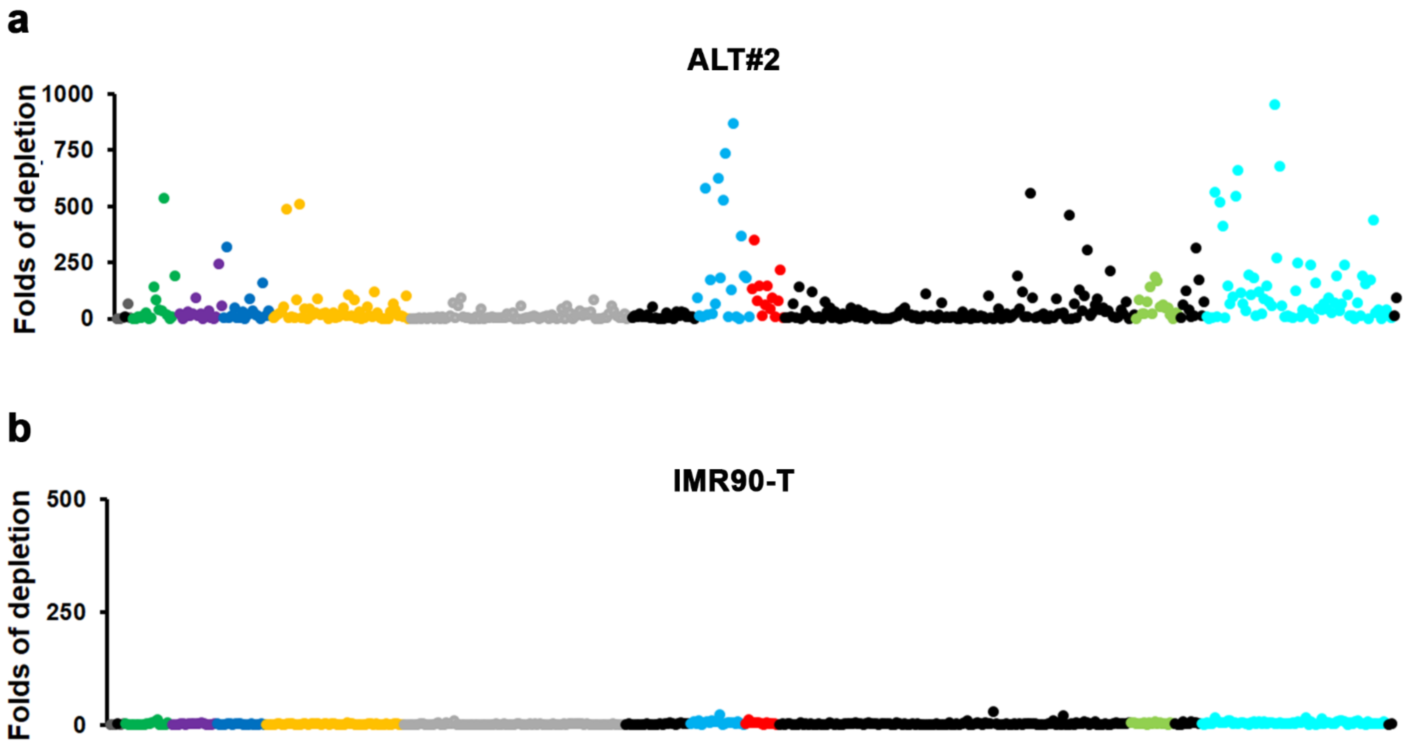


**Supplementary Figure 5. CRISPR-based KDM2A tiling assay. a, b** CRISPR-based KDM2A tiling assay in ALT#2 (**a**) or IMR90-T cells (**b**). Plotted is the fold changes of sgRNA abundance (ratio of start to end point) after 16 population doublings in culture. x axis shows targeting sgRNAs; the domain location of each sgRNA within KDM2A protein is indicated by colors. y axis shows the fold changes of individual KDM2A targeting sgRNAs following the culture period.

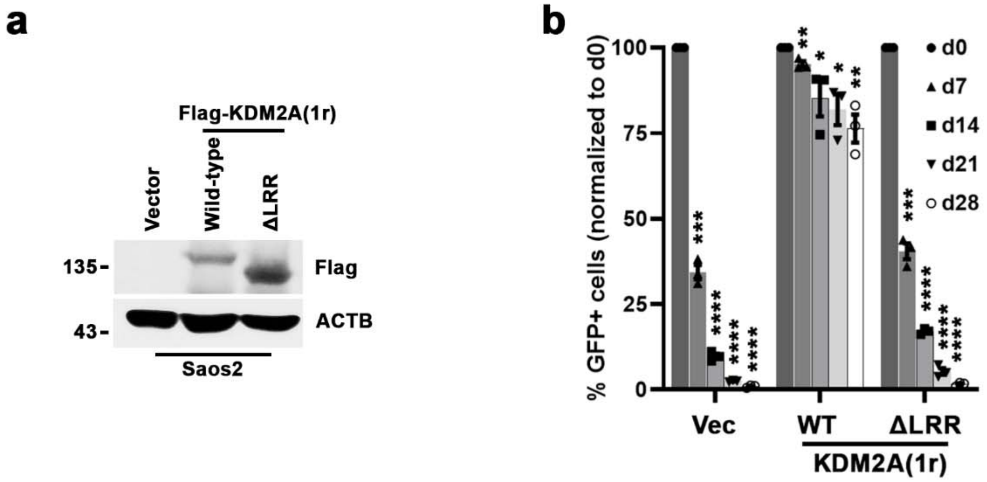


**Supplementary Figure 6. LRR domain of KDM2A is required for its ALT supporting function. a** Western blot analysis of Flag and ACTB (loading control) in whole-cell lysates prepared from Saos2 cells transduced with construct encoding KDM2A wild-type (aa 1-aa 1161) or ΔLRR mutant (aa 1-aa 945). The Flag-tagged wild-type and ΔLRR mutant were generated from CRISPR-resistant KDM2A-1r construct. **b** Competition-based proliferation assay of GFP-coexpressing sgK#1 in Saos2 cells transduced with KDM2A wild-type or ΔLRR mutant construct. The bar graphs are expressed as means ± s.e.m. of three independent experiments; paired t-test. * denotes ns (not significant); ** $P < 0.05$; *** $P < 0.01$; **** $P < 0.001$.

Supplementary Figure 7

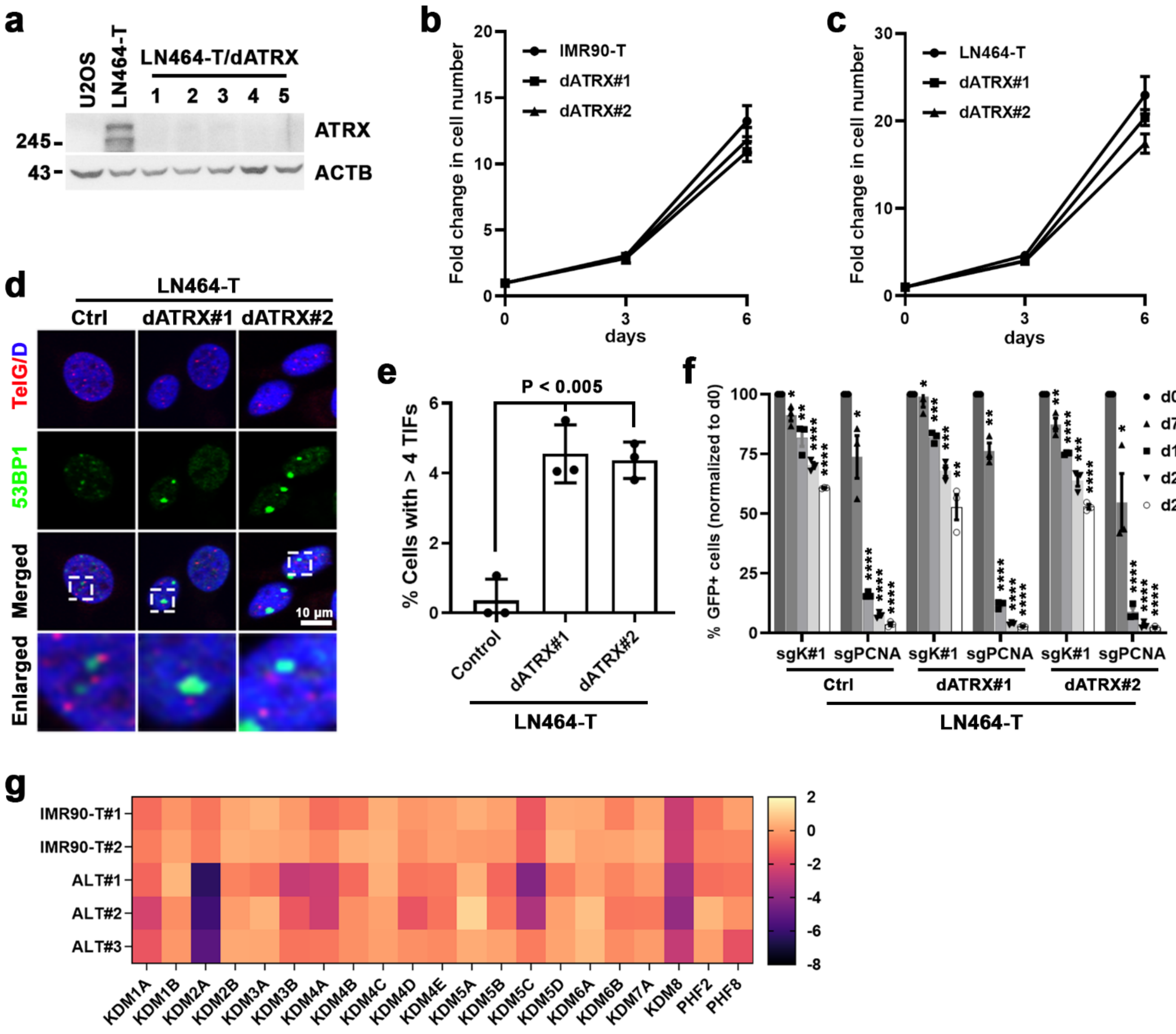


**Supplementary Figure 7. KDM2A is not essential for non-ALT ATRX-null cells. a** Western blot analysis of ATRX expression in whole-cell lysates prepared from sgCtrl or ATRX-depleted LN464-T cells. The ATRX depleted lines (dATRX clone#1-5) were established from clonally isolated LN464-T cells transduced with ATRX-targeted sgRNA. The lysates of U2OS and LN464-T cells were included as a negative or a positive control of ATRX expression respectively. **b** Cell proliferation of wild-type or ATRX-depleted (dATRX#1 and dATRX#2) IMR90-T cells. **c** Cell proliferation of wild-type or ATRX-depleted (dATRX#1 and dATRX#2) LN464-T cells. **d** Representative immuno-FISH images of 53BP1 and telomeres in control or ATRX-depleted LN464-T cells (dATRX#1 and dATRX#2). Scale bar, 10 µm. **e** Percentages of cells containing ≥ 4 53BP1-associated telomere dysfunction-induced foci (TIFs). Data are expressed as means ± s.e.m. of three independent experiments; unpaired t-test. **f** Competition-based proliferation assay of KDM2A-targeted sgK#1 (linked with GFP expression) in control or ATRX-depleted LN464-T cell lines. Data are expressed as means ± s.e.m. of three independent experiments; paired t-test. * denotes ns (not significant); ** $P < 0.05$; *** $P < 0.01$; **** $P < 0.001$. **g** Heatmap depicting the gene dependency score (GDS) of the indicated histone demethylases in IMR90-T (T#1 and T#2), ALT#1, ALT#2, or ALT#3 cells. The GDS score was calculated by averaging the log2FC of all sgRNAs targeting that gene after 16 population doubling.

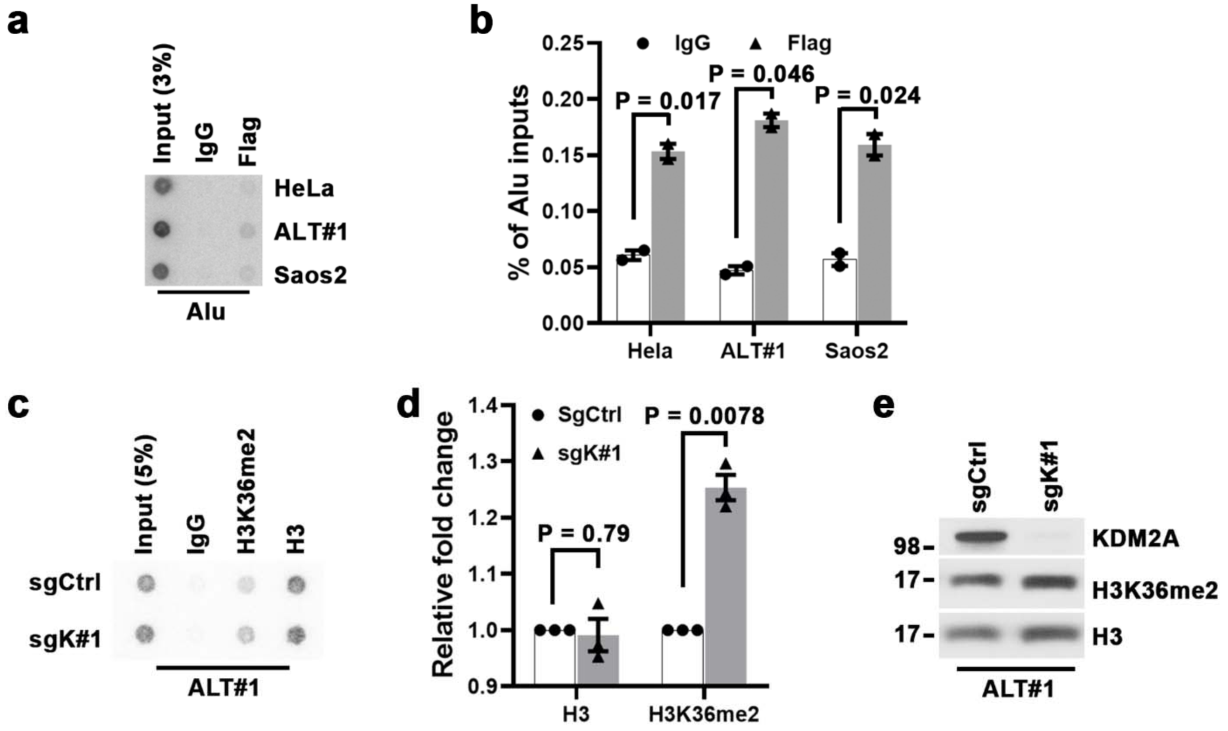


**Supplementary Figure 8. Depletion of KDM2A in ALT cells elevates telomere H3K36me2.** **a, b** Alu dot blot analysis (**a**) and quantification (**b**) of anti-Flag or IgG chromatin immune-precipitation (ChIP) in the indicated cell lines. The IgG ChIP was included as a control for non-specific signal. The input and ChIP DNAs processed against the indicated antibodies were assayed by dot-blotting and hybridized with $^{32}$P-labeled Alu probe. The relative enrichment was calculated after normalization of ChIP DNA signals to the respective input DNA signals. Data are expressed as means ± s.e.m. of two independent experiments; paired t-test. **c, d** Telomere dot blot analysis (**c**) and quantification (**d**) of anti-H3K36me2, anti-H3 or IgG ChIP in sgCtrl or sgK#1-transduced ALT#1 cells. The relative enrichment was calculated after normalization to the respective ChIP DNA signals of sgCtrl-transduced samples. Notes, the H3K36me2 ChIP DNA signals were first normalized to their respective H3 ChIP signals. Data are expressed as means ± s.e.m. of three independent experiments; paired t-test. **e** Western blot analysis of KDM2A, H3K36me2, and H3 in lysates prepared from sgCtrl or sgK#1-transduced ALT#1 cells.

Supplementary Figure 9

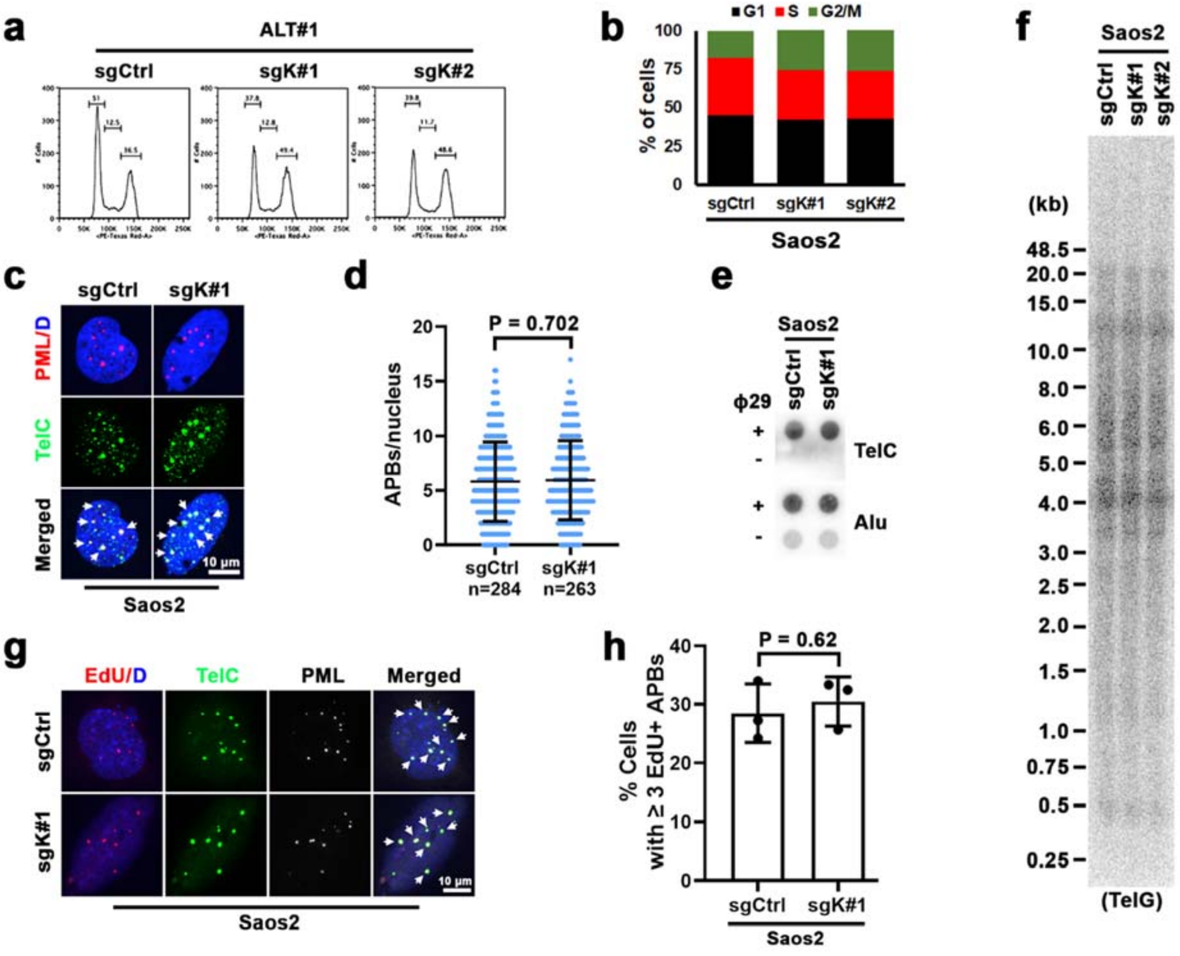


**Supplementary Figure 9. KDM2A depletion induces G2/M accumulation without affecting ALT-associated activities. a** Cell cycle phase analysis of sgCtrl, sgK#1 or sgK#2-transduced ALT#1 cells. **b** Quantification of cell cycle distribution of sgCtrl, sgK#1 or sgK#2-transduced Saos2 cells. **c** Representative immuno-FISH images of ALT-associated PML bodies (APBs) in sgCtrl or sgK#1-transduced Saos2 cells. The arrows denote APB foci. Scale bar = 10 µm. **d** Quantifications of APB formation as in (**c**). Each dot represents an individual nucleus. Means ± s.e.m. of three independent experiments; unpaired t-test. **e** C-circle assays of Saos2 cells transduced with sgCtrl or sgK#1. Genomic DNAs were prepared from the indicated cells at day 10 post sgRNA transduction and assayed by $^{32}$P-labeled TelG probe. **f** Telomere restriction fragment analysis of telomere length in sgCtrl, sgK#1 or sgK#2-transduced Saos2 cells. Genomic DNAs prepared from the indicated cells at day 10 post lentiviral sgRNA infection were assayed by $^{32}$P-labeled TelG probe. **g** Representative immuno-FISH images of EdU colocalized APBs (EdU-APBs) in sgCtrl or sgK#1-transduced Saos2 cells. Cells were synchronized in G2 with sequential thymidine and CDK1 inhibitor treatment before EdU labeling for 2 h. EdU was assayed by Click-It reaction, PML was analyzed by IF, and telomeres were detected by FISH. The arrows denote EdU-APB foci. Scale bar, 10 μm. **h** Percentages of cells containing ≥ 3 EdU-APB foci are expressed as means ± s.e.m. of three independent experiments; unpaired t-test.

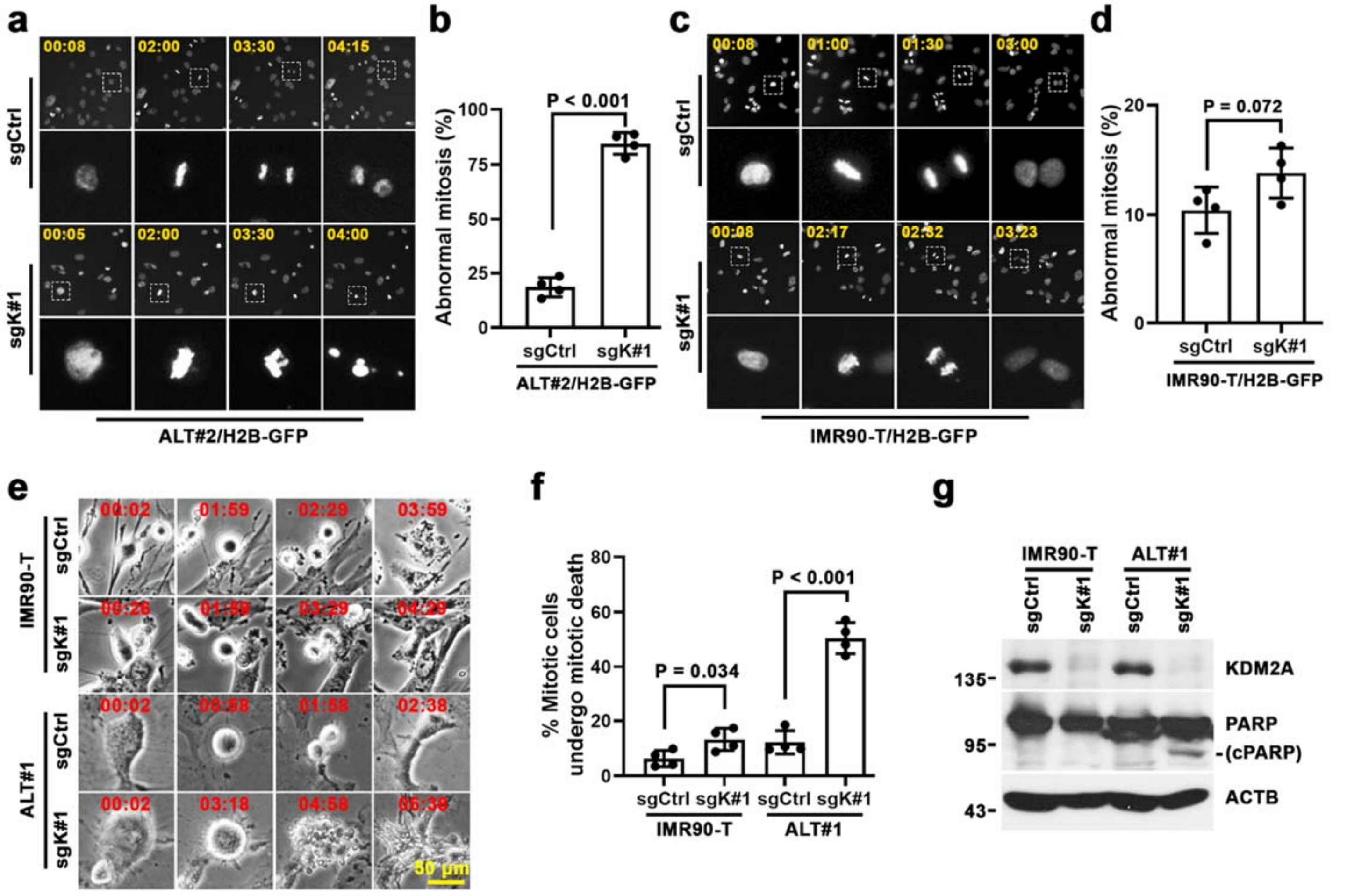


**Supplementary Figure 10. KDM2A depletion induces mitotic death of ALT cells. a** Representative frames of time-lapse fluorescence live cell imaging of GFP-H2B-expressing ALT#2 cells transduced with sgCtrl or sgK#1. Cells were synchronized in G2 with sequential thymidine and CDK1 inhibitor treatment before timed release into M phase. Time is shown as (hours: minutes) relative to the first image of the series. **b** Percentages of aberrant mitosis are expressed as means ± s.e.m. of four independent experiments; unpaired t-test. **c** Representative time-lapse fluorescence live cell imaging frames of GFP-H2B-expressing IMR90-T cells transduced with sgCtrl or sgK#1. Cells were synchronized in G2 before timed release into M phase. **d** Percentages of aberrant mitosis are expressed as means ± s.e.m. of four independent experiments; unpaired t-test. **e** Representative images from live cell imaging experiments depicting mitotic outcome of IMR90-T and ALT#1 cells transduced with sgCtrl or sgK#1. **f** Percentages of mitotic cells that underwent mitotic death are expressed as means ± s.e.m. of four independent experiments; unpaired t-test. **g** Western blot analysis of KDM2A, PARP1 and ACTB in whole-cell lysates prepared from sgCtrl or sgK#1-transduced IMR90-T or ALT#1 cells. cPARP denotes cleaved PARP1.

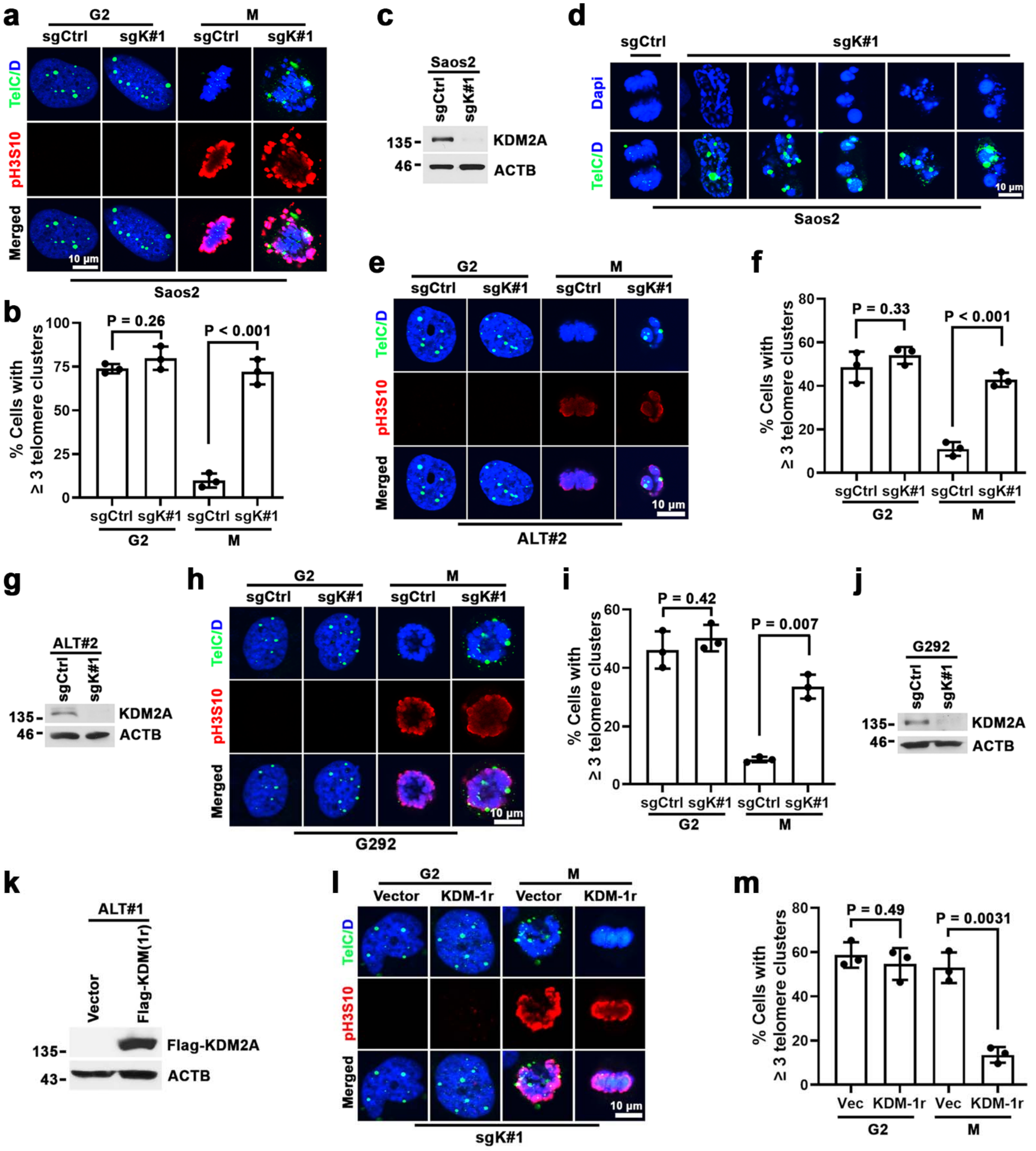


**Supplementary Figure 11. KDM2A depletion impairs ALT telomere de-clustering. a** Representative immuno-FISH images of telomere clusters in sgCtrl or sgK#1-transduced G2 or histone H3-Ser10 (pH3S10)+ mitotic Saos2 cells. The G2 cells were synchronized from sequential thymidine and CDK1 inhibitor treatment. The mitotic (M) cells were from G2 synchronized cells upon 4 h release from CDK1 inhibitor. Scale bar, 10 μm. **b** Percentages of cells containing ≥ 3 clustered telomere foci. Data are expressed as means ± s.e.m. of three independent experiments; unpaired t-test. **c** Western blot analysis of KDM2A and ACTB in whole-cell lysates prepared from sgCtrl or sgK#1-transduced Saos2 cells. **d** Representative images of abnormal mitotic telomere clusters in sgK#1-transduced Saos2 cells. G2 synchronized sgCtrl or sgK#1-transduced Saos2 cells were released into mitosis for 6 h. Telomeres were detected by FISH. **e** Representative immuno-FISH images of telomere clusters in sgCtrl or sgK#1-transduced G2 synchronized or histone H3-Ser10 (pH3S10)+ mitotic ALT#2 cells. The M-phased cells were from G2 synchronized cells upon 2 h release from CDK1 inhibitor. Scale bar, 10 μm. **f** Percentages of the indicated G2 or M-phased ALT#2 cells containing ≥ 3 clustered telomere foci. Data are expressed as means ± s.e.m. of three independent experiments; unpaired t-test. **g** Western blot analysis of KDM2A and ACTB in whole-cell lysates prepared from sgCtrl or sgK#1-transduced ALT#2 cells. **h** Representative immuno-FISH images of telomere clusters in sgCtrl or sgK#1-transduced G2 synchronized or mitotic G292 cells. The M-phased cells were from G2 synchronized G292 cells upon 4 h release from CDK1 inhibitor. Scale bar, 10 μm. **i** Percentages of the indicated G2 or M-phased G292 cells containing ≥ 3 clustered telomere foci. Data are expressed as means ± s.e.m. of three independent experiments; unpaired t-test. **j** Western blot analysis of KDM2A and ACTB in whole-cell lysates prepared from sgCtrl or sgK#1-transduced G292 cells. **k** Western blot analysis of Flag and ACTB in whole-cell lysates prepared from ALT#1 cells transduced with vector control or sgK#1-resistance Flag-KDM2A(r). **l** Representative immuno-FISH images of telomere clusters in sgK#1-transduced G2 or mitotic ALT#1 cells complemented with vector control or Flag-KDM2A(r). Scale bar, 10 μm. **m** Percentages of the indicated G2 or M-phased ALT#1 cells containing ≥ 3 clustered telomere foci. Data are expressed as means ± s.e.m. of three independent experiments; unpaired t-test.

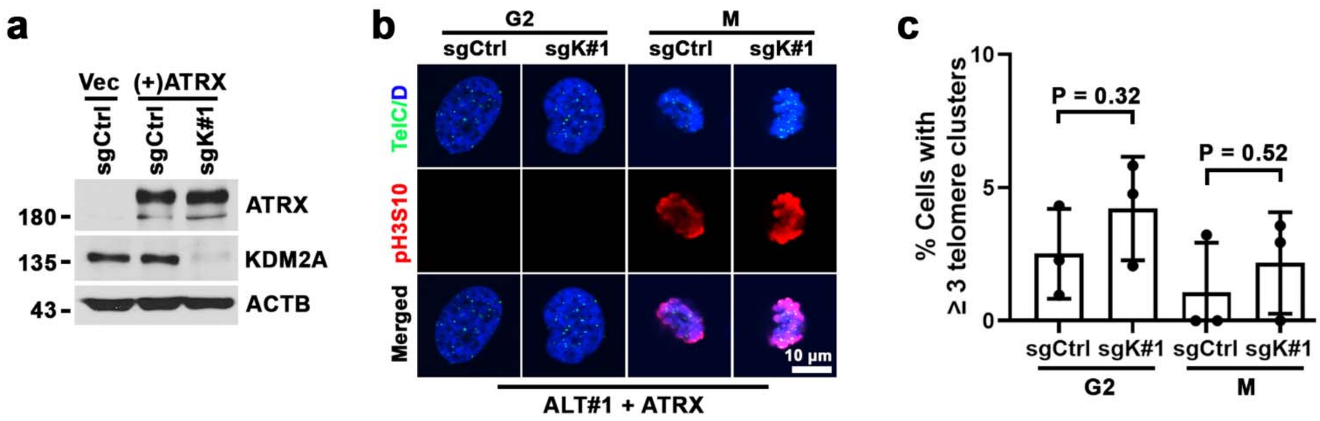


**Supplementary Figure 12. KDM2A is not required for mitotic telomere segregation of ALT#1 cells complemented with ATRX. a** Western blot analysis of ATRX, KDM2A and ACTB in whole-cell lysates prepared from sgCtrl or sgK#1-transduced ALT#1 cells complemented with ATRX cDNA. **b** Representative immuno-FISH images of telomere clusters in sgCtrl or sgK#1-transduced G2 and mitotic ALT#1 cells complemented with *ATRX* cDNA. Scale bar, 10 µm. **c** Percentages of the indicated G2 or M-phased ALT#1 cells containing ≥ 3 clustered telomere foci. Data are expressed as means ± s.e.m. of three independent experiments; unpaired t-test.

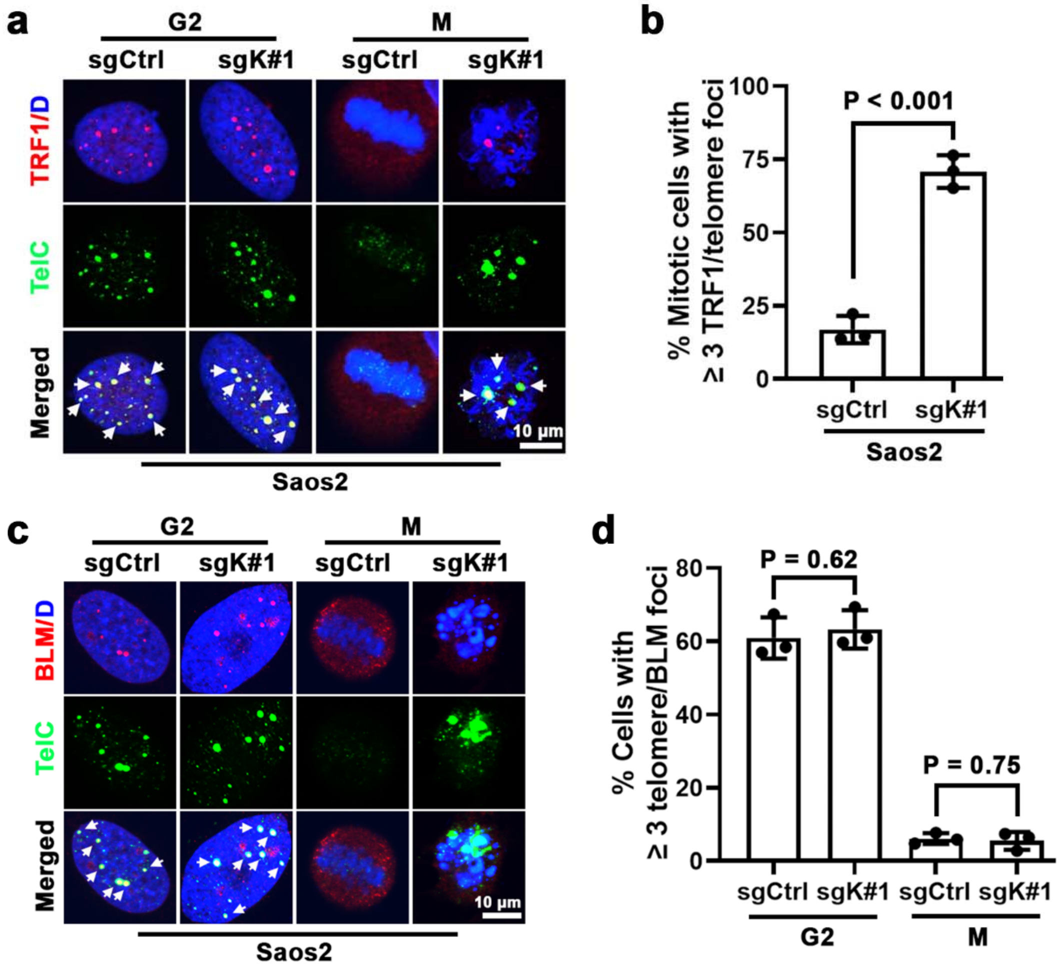


**Supplementary Figure 13. Depletion of KDM2A does not affect BLM recruitment to ALT telomeres. a** Representative immuno-FISH images of TRF1-telomere association in G2 or M-phase of sgCtrl or sgK#1-transduced Saos2 cells. The M-phase cells were from G2 synchronized cells upon 4 h release from CDK1 inhibitor. The arrows denote TRF1-associated telomere foci. **b** Percentages of mitotic cells containing ≥ 3 TRF1-associated telomere foci. Data are expressed as means ± s.e.m. of three independent experiments; unpaired t-test. **c** Immuno-FISH analysis of BLM and telomeres colocalization in G2- or M-phase Saos2 cells transduced with sgCtrl or sgK#1. The arrows denote BLM-associated telomere foci. **d** Percentages of cells containing ≥ 3 BLM-associated telomere foci. Data are expressed as means ± s.e.m. of three independent experiments; unpaired t-test. Scale bar, 10 µm.

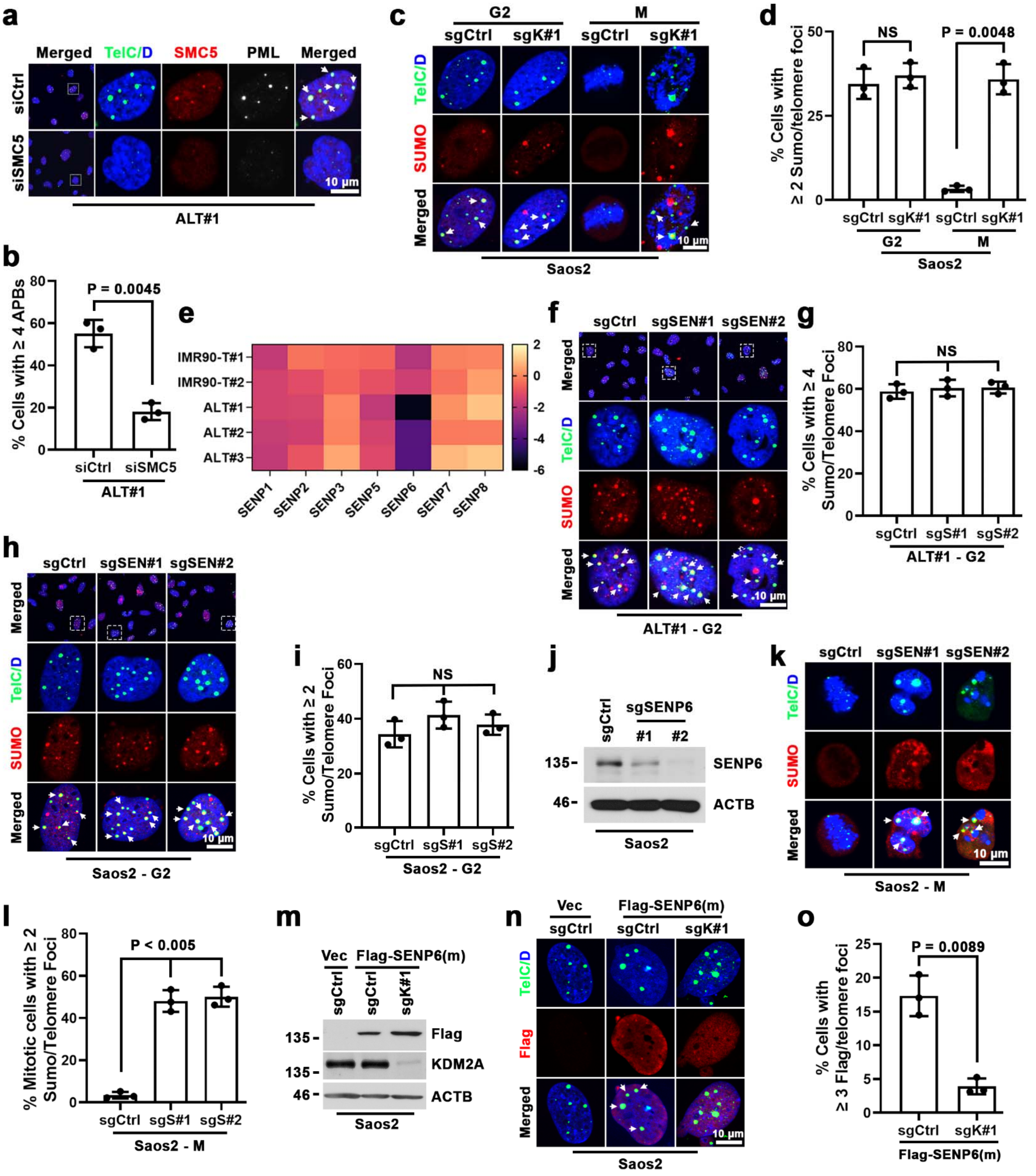


**Supplementary Figure 14. KDM2A depletion impairs ALT telomere de-SUMOylation. a** Representative immuno-FISH images of APBs in ALT#1 cells transfected with control or SMC5 siRNA. The arrows denote APB foci. Scale bar, 10 µm. **b** Percentages of cells containing ≥ 4

APB foci are expressed as means ± s.e.m. of three independent experiments; unpaired t-test. **c** Representative immuno-FISH images of telomere SUMOylation in sgCtrl or sgK#1-transduced Saos2 cells. The mitotic (M) cells were from G2 synchronized cells upon 4 h release from CDK1 inhibitor. The arrows denote SUMO2/3-associated telomere foci. **d** Percentages of cells containing ≥ 2 SUMO-associated telomere foci. Data are expressed as means ± s.e.m. of three independent experiments; unpaired t-test. **e** Heatmap depicting the gene dependency score (GDS) of the indicated SENPs in IMR90-T (T#1 and T#2), ALT#1, ALT#2, or ALT#3 cells. The GDS score was calculated by averaging the log2FC of all sgRNAs targeting that gene after 16 population doubling. **f** Immuno-FISH analysis of telomere SUMOylation in sgCtrl, sgSEN#1 or sgSEN#2-transduced ALT#1 cells in G2-phase. The G2 cells were synchronized from sequential thymidine and CDK1 inhibitor treatment. The arrows denote SUMO2/3-associated telomere foci. **g** Percentages of G2 cells containing ≥ 4 SUMO-associated telomere foci are expressed as means ± s.e.m. of three independent experiments; unpaired t-test. **h** Immuno-FISH analysis of telomere SUMOylation in sgCtrl, sgSEN#1 or sgSEN#2-transduced Saos2 cells in G2-phase. The arrows denote SUMO2/3-associated telomere foci. **i** Percentages of the indicated Saos2 cells containing ≥ 2 SUMO-associated telomere foci. Data are expressed as means ± s.e.m. of three independent experiments; unpaired t-test. **j** Western blot analysis of SENP6 expression in whole-cell lysates prepared from sgCtrl, sgSEN#1 or sgSEN#2-transduced Saos2 cells. **k** Immuno-FISH analysis of telomere-associated SUMOylation in mitotic Saos2 cells transduced with sgCtrl, sgSEN#1 or sgSEN#2. The mitotic cells were from G2 synchronized cells after 4 h release from CDK1 inhibitor. **l** Percentages of the indicated mitotic cells containing ≥ 2 SUMO2/3-associated telomere foci. Data are expressed as means ± s.e.m. of three independent experiments; unpaired t-test. **m** Western blot analysis of Flag, KDM2A and ACTB in whole-cell lysates prepared from the indicated Saos2 cells transduced with vector control or Flag-tagged SENP6C1030A mutant. **n** Immuno-FISH analysis of telomere-SENP6$^{C1030A}$ association in sgCtrl or sgK#1-transduced Saos2 cells infected with lentiviral Flag-tagged SENP6$^{C1030A}$ mutant. The arrows denote SENP6$^{C1030A}$-associated telomere foci. **o** Percentages of cells containing ≥ 3 Flag-SENP6$^{C1030A}$-associated telomere foci. Data are expressed as means ± s.e.m. of three independent experiments; unpaired t-test.

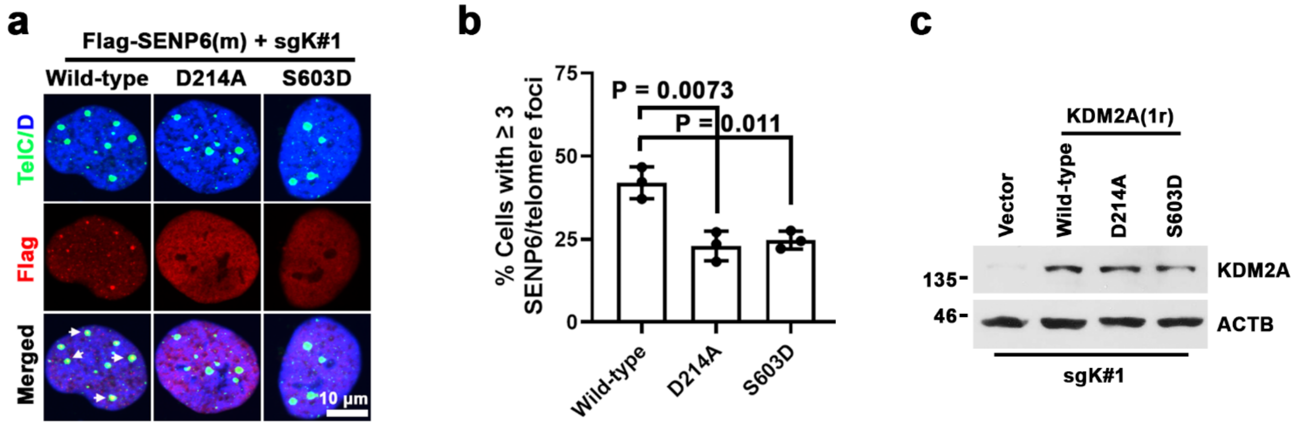


**Supplementary Figure 15. KDM2A facilitates SENP6 recruitment to ALT telomeres. a** Immuno-FISH analysis of telomere-SENP6$^{C1030A}$ association in sgK#1 and Flag-tagged SENP6$^{C1030A}$-transduced ALT#1 cells complemented with sgK#1-resistant wild-type or indicated KDM2A mutant. The arrows denote SENP6$^{C1030A}$-associated telomere foci. **b** Percentages of cells containing ≥ 3 Flag-SENP6$^{C1030A}$-associated telomere foci. Data are expressed as means ± s.e.m. of three independent experiments; unpaired t-test. **c** Western blot analysis of KDM2A and ACTB in whole-cell lysates prepared from sgK#1 and Flag-tagged SENP6$^{C1030A}$-transduced ALT#1 cells that were complemented with the indicated wild-type or KDM2A mutant.

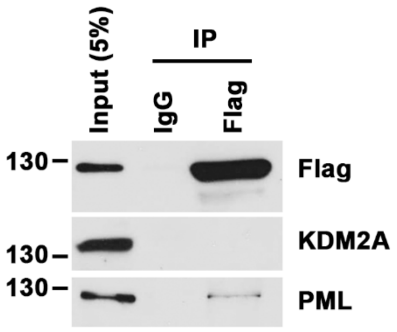


**Supplementary Figure 16. SENP6 does not directly bind to KDM2A.** Western blot analysis of KDM2A and PML in anti-Flag or IgG immune-precipitation (IP) of whole-cell lysates prepared from Flag-tagged SENP6$^{C1030A}$-transduced ALT#1 cells.
